# Rescue of FTLD-associated TDP-43 pathology and neurodegeneration by peripheral AAV-mediated expression of brain-penetrant progranulin

**DOI:** 10.1101/2023.07.14.549089

**Authors:** Marvin Reich, Matthew J. Simon, Beate Polke, Georg Werner, Christian Schrader, Iñaki Paris, Sophie Robinson, Sonnet S. Davis, Gabrielly Lunkes de Melo, Lennart Schlaphoff, Lena Spieth, Stefan Berghoff, Todd Logan, Brigitte Nuscher, Katrin Buschmann, Dieter Edbauer, Mikael Simons, Jung H. Suh, Thomas Sandmann, Mihalis S. Kariolis, Sarah L. DeVos, Joseph W. Lewcock, Dominik Paquet, Anja Capell, Gilbert Di Paolo, Christian Haass

## Abstract

Progranulin (PGRN) haploinsufficiency is a major risk factor for frontotemporal lobar degeneration with TDP-43 pathology (FTLD-*GRN*). Multiple therapeutic strategies are in clinical development to restore PGRN levels in the CNS, including gene therapy. However, a limitation of current gene therapy approaches aimed to alleviate FTLD-associated pathologies may be their inefficient brain exposure and biodistribution. We therefore developed an adeno-associated virus (AAV) targeting the liver (L) to achieve sustained peripheral expression of a transferrin receptor (TfR) binding, brain-penetrant (b) PGRN variant (AAV(L):bPGRN) in two mouse models of FTLD-*GRN*, namely *Grn* knockout and *GrnxTmem106b* double knockout mice. This therapeutic strategy avoids potential safety and biodistribution issues of CNS-administered AAVs while maintaining sustained levels of PGRN in the brain following a single dose. AAV(L):bPGRN treatment reduced several FTLD-*GRN* associated disease pathologies including severe motor function deficits, aberrant TDP-43 solubility and phosphorylation, dysfunctional protein degradation, lipid metabolism, gliosis and neurodegeneration in the brain. Translatability of our findings was confirmed in a novel human *in vitro* model using co-cultured human induced pluripotent stem cell (hiPSC)-derived microglia lacking PGRN and TMEM106B and wild-type hiPSC-derived neurons. As in mice, aberrant TDP-43, lysosomal dysfunction and neuronal loss were ameliorated after treatment with exogenous TfR-binding protein transport vehicle fused to PGRN (PTV:PGRN). Together, our studies suggest that peripherally administered brain-penetrant PGRN replacement strategies can ameliorate FTLD-*GRN* relevant phenotypes including TDP-43 pathology, neurodegeneration and behavioral deficits. Our data provide preclinical proof of concept for the use of this AAV platform for treatment of FTLD-*GRN* and potentially other CNS disorders.

**One sentence summary:** Peripheral AAV-mediated delivery of brain-penetrant PGRN rescues TDP-43 pathology, neurodegeneration and motor phenotypes in FTLD-*GRN* models.

## INTRODUCTION

Haploinsufficiency of progranulin (PGRN) caused by heterozygous loss-of-function (LoF) mutations in the *GRN* gene dramatically enhances the risk for FTLD-TDP (*1–3*). As in all TDP-43 proteinopathies, FTLD with PGRN deficiency (FTLD-*GRN*) displays aggregation, pathological processing and abnormal phosphorylation of TDP-43 (*4, 5*). FTLD-*GRN* patients present with severe clinical symptoms associated with underlying neurodegeneration primarily in the cortex (*5, 6*), resulting in enhanced neurofilament light chain (NfL) levels in blood and cerebrospinal fluid (CSF) (*7–9*), as well as pronounced neuroinflammation and dysfunction of the endo-lysosomal system (*5, 10, 11*). PGRN deficiency is hypothesized to contribute to these pathologies through its established role in lysosomal function (*11–14*), microglia homeostasis (*15–17*) and inflammatory signaling (*11, 18, 19*) (reviewed in (*4, 5, 20, 21*)). PGRN or its proteolytic derivatives, the granulin peptides, are localized within the endo-lysosomal system (*22, 23*) and are implicated in the regulation of lysosomal proteases (*11, 12, 24–27*), lysosomal pH (*14, 28, 29*) as well as lipid hydrolases, such as glucocerebrosidase (GCase) (*29–34*). Loss of PGRN in mice causes an age-dependent increase in lysosomal enzymes, lysosomal membrane proteins, hyperactivated microglia and saposin D, indicating deficits in endolysosomal and autophagic protein degradation (*11–19, 25, 29–33, 35, 36*). Lysosomal phenotypes further include deficiency of endolysosomal anionic phospholipid bis(monoacylglycero)phosphate (BMP) and secondary storage of sphingolipids, such as GCase substrates and gangliosides (*29, 33, 37*). Consistent with its critical role in the lysosome, homozygous LoF mutations in *GRN* cause the fatal lysosomal storage disease (LSD) termed neuronal ceroid lipofuscinosis (NCL) in humans (*38, 39*).

*Grn^+/–^* (*Grn* HET) mice present with no robust disease-relevant phenotypes (*12, 40*). In contrast, *Grn^−/−^*(*Grn* KO) mice recapitulate lysosomal and inflammatory FTLD-*GRN* pathologies but exhibit only minor TDP-43 pathology even at advanced ages (*12, 16, 19*). Furthermore, behavioral phenotypes in *Grn* HET and *Grn* KO mice are subtle and have limited relevance to FTLD-*GRN* clinical phenotypes (*40, 41*). We and others recently developed a mouse model that addressed some of the limitations of the *Grn* KO model by introducing an additional KO of *Tmem106b* (*42–44*), a genetic risk factor for FTLD-*GRN* (*45*). TMEM106B is involved in endo-lysosomal degradation and trafficking (*46–50*) and has been shown to undergo processing and give rise to a C-terminal fragment prone to forming fibrils which are observed in the brain of FTLD-*GRN* patients and aged individuals (*51–53*). The *Tmem106b* and *Grn* double KO (DKO) generally exacerbates phenotypes observed in the single KOs, including lysosomal and autophagic dysfunction, neuroinflammation, and myelin loss (*42–44, 54*). Importantly, unlike the *Grn* KO, it exhibits much more robust accumulation of insoluble and hyperphosphorylated TDP-43, and motor impairment at an age of 4 months (*42–44*). Thus, this model reproduces key biochemical, cell biological as well as disease-relevant motor and neurodegeneration phenotypes of FTLD-*GRN*.

To date, there is no disease-modifying treatment for FTLD-*GRN*, although several therapeutic approaches are currently being tested in the clinic (reviewed in (*20, 55*)). Given that it is a monogenic disease, correcting the PGRN deficiency with a protein replacement therapy may offer an effective therapeutic approach to slow or halt disease progression and reduce pathology. While adeno-associated virus (AAV)-based therapies are under clinical investigation (*56–58*), they typically rely on direct CNS administration, which may introduce safety concerns as well as result in limited or uneven biodistribution of the AAV (*56–58*) (reviewed in (*59*)). Thus, the number of cells expressing PGRN and PGRN exposure in cells far from the injection site might be insufficient. Consistent with this, a recent study suggests that AAV1 and 9 administered to the CSF can reduce disease phenotypes in the spinal cord, but not in the brain of an FTLD-*GRN* mouse model, limiting their use for neurodegenerative diseases with widespread brain pathologies, such as FTLD-*GRN* (*60*). As an alternative, we recently fused human recombinant PGRN to a transport vehicle (PTV:PGRN), enabling transferrin receptor (TfR)-mediated transcytosis across the blood-brain barrier (BBB) and delivery throughout the whole CNS (*29*). Using PTV:PGRN, multiple FTLD-related pathologies in *Grn* KO mice and human microglia – lipid/lysosomal abnormalities, microglial/astrocytic hyperactivation – were rescued (*29*). However, due to the limited pathology of the *Grn* KO mouse model, TDP-43 pathology, motor deficits and neurodegeneration could not be investigated.

Here, we developed an advanced gene therapy approach by using a liver-targeting (L) adeno-associated virus expressing brain-penetrant (b) PGRN (AAV(L):bPGRN). This strategy combines a TfR-mediated, brain-penetrant PGRN biologic with AAV-targeted delivery to the liver, enabling brain-wide PGRN delivery with a single peripheral administration treatment paradigm without requiring direct CNS injection. We tested the AAV(L):bPGRN approach in *Grn* KO mice as well as in our recently described DKO mouse model lacking both *Grn* and *Tmem106b* to study whether PGRN replacement could rescue TDP-43 pathology, motor impairment, exacerbated neuroinflammation, neuronal loss, impaired protein degradation, and lipidomic and transcriptomic changes in the brain. To complement our *in vivo* efforts, we generated a hiPSC-derived FTLD-model by co-culturing *GRN* and *TMEM106B* DKO human iPSC-derived microglia (iMG) with WT human iPSC-derived neurons (iN). This model recapitulates multiple *in vivo* phenotypes, including microglial hyperactivation, lysosomal abnormalities, aberrant TDP-43 processing and phosphorylation, as well as neurodegeneration. Strikingly, our replacement strategy significantly reduced all phenotypes observed in the DKO mice and human iPSC model, despite the phenotypic severity and constitutive loss of TMEM106B.

Our study provides compelling evidence that brain-penetrant PGRN replacement therapies such as AAV(L):bPGRN or PTV:PGRN can have an impact on the central pathologies of FTLD-*GRN* in the brain, namely TDP-43 pathology and neurodegeneration, in both mouse and human FTLD models.

## RESULTS

### AAV(L):bPGRN allows safe and robust expression and brain delivery

We generated AAV(L):bPGRN, a liver-targeting AAV8 encoding a fusion protein (8D3:PGRN) consisting of a single chain fragment variable (scFv) antibody recognizing mouse transferrin receptor (TfR; 8D3) fused to human PGRN (hPGRN). AAV(L):bPGRN infects hepatocytes in the liver where it stably expresses 8D3:PGRN after a single peripheral administration to provide sustained expression and secretion of the fusion protein into systemic circulation (Fig. 1A). As a proof-of-principle, we first confirmed AAV(L):bPGRN treatment drives stable expression of 8D3:PGRN in *Grn* KO mice. We infected 4-5-month-old WT and *Grn* KO mice with AAV(L):bPGRN and monitored hPGRN levels in liver, brain and plasma for 31 weeks (Fig. 1B). In the liver of AAV(L):bPGRN treated mice, hPGRN levels of 5 µg/mg above background were observed 9 weeks after dosing, suggesting robust liver targeting of AAV(L):bPGRN (Fig. 1C). Furthermore, one week after injection, hPGRN was detectable in plasma and after 4 weeks, expression stabilized at an average of 3 µg/ml (Fig. 1D), demonstrating that the liver expressed AAV(L):bPGRN is secreted into circulation. Finally, robust levels of hPGRN were detected in the brain of mice at both 9 and 31 weeks after injection, supporting efficient blood-brain-barrier transcytosis of the fusion protein (Fig. 1E).

**Fig. 1:**
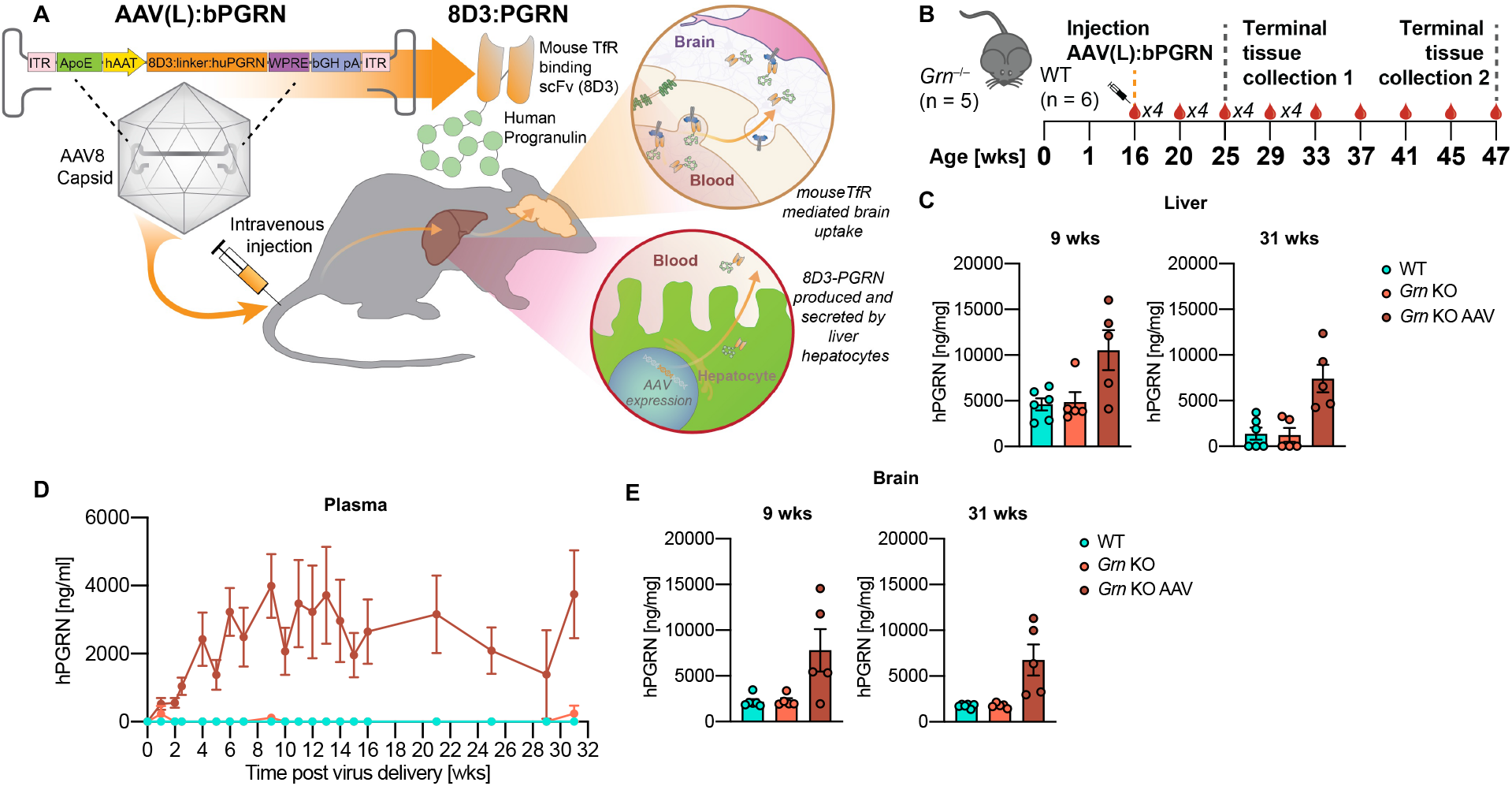
AAV-expressed PGRN is delivered to the brain and shows no side effects in *Grn* KO mice. (**A**) Graphical summary of treatment paradigm. Intravenously administered AAV expresses brain-penetrant (b) human progranulin (PGRN) in the liver (L) (AAV(L):bPGRN). PGRN is fused to a single chain fragment variable (scFv) antibody recognizing mouse transferrin receptor (TfR; 8D3). 8D3:PGRN is expressed by liver hepatocytes, released into the vasculature and can cross the blood-brain-barrier *via* transferrin receptor-binding. (**B**) Schematic of AAV treatment and sample collection in the *Grn* KO AAV treatment study. Directly after treatment, blood was sampled weekly (x4 = 4 times in 4 weeks (wks)), from 16 weeks after treatment on every 4 weeks. (**C-E**) hPGRN levels in liver (C), plasma (D) and brain (E) following AAV(L):bPGRN treatment in mice. For tissues, 9-week samples are depicted on the left and 31-week on the right. For all figures, error bars represent standard error of the mean (SEM). n=5-6/group.

As high levels of AAV infection have been linked to adverse effects, including hepatotoxicity (*61*), and TfR is highly expressed on immature red blood cells (RBCs) we evaluated serum chemistry (including alanine transaminase (ALT) and aspartate aminotransferase (AST)) and hematology (including circulating reticulocytes), and performed histopathologic analysis of key organs from *Grn* KO mice treated with AAV(L):bPGRN after 31 weeks (Fig. S1, Table S1). No significant microscopic findings were noted in the assessed tissues, including brain, liver, heart, and sciatic nerve (Table S1). No treatment-related findings were identified in serum. Minimal and non-adverse changes were observed in hematology, including slight decrease in red blood cell size (mean corpuscular volume (MCV)) and hemoglobin (mean corpuscular hemoglobin (MCH)) levels, accompanied by a slight increase in RBC and reticulocyte counts (Fig. S1A-D). Thus, a single injection of AAV(L):bPGRN in mice allows for sustained brain delivery of hPGRN without any significant deleterious side-effects.

### AAV(L):bPGRN is stably expressed in DKO mice and ameliorates motor defects

We next examined the effects of AAV(L):bPGRN mediated PGRN replacement in DKO mice. Previously, we demonstrated that DKO mice not only exacerbate the FTLD-related pathologies observed in *Grn* KO mice, but additionally exhibit motor impairment, and pronounced abnormal phosphorylation and deposition of insoluble TDP-43 (*42–44*). To determine whether restoring PGRN is sufficient to prevent these phenotypes, DKO mice were injected at 6 weeks of age, prior to development of robust pathologies. A predetermined treatment duration until an age of 15 weeks was established due to the severity of the DKO phenotypes (Fig. 2A) (*42*). As early as 4 weeks after AAV delivery, stable levels of hPGRN of around 4 µg/ml were observed in plasma of treated WT and DKO mice (Fig. 2B). Robust hPGRN protein levels were also detected in liver and brain of treated mice (Fig. 2C,D), without affecting endogenous expression of mPGRN (Fig. S2A-C).

**Fig. 2:**
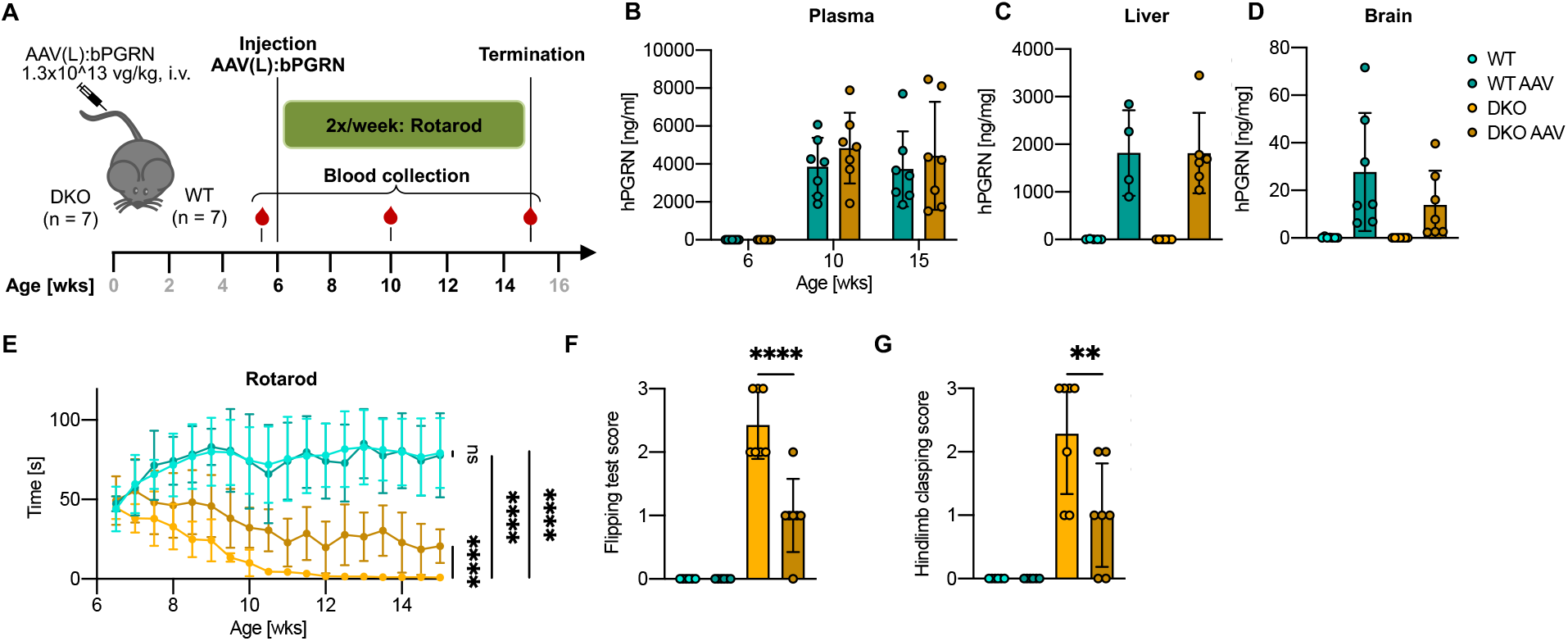
AAV-expressed PGRN is delivered to the brain and ameliorates behavioral abnormalities in DKO mice. (**A**) Schematics of AAV treatment and sample collection in the *Grn* x *Tmem106b* double knockout (DKO) AAV(L):bPGRN treatment study. (**B-D**) hPGRN levels in plasma (left, n=7/group; 6, 10 and 15-week-old), liver (middle, n=4-6/group; 15-week-old) and brain (right, n=7/group; 15-week-old) following AAV(L):bPGRN treatment in mice as assessed by ELISA. (**E**) Longitudinal assessment of rotarod performance following treatment. (**F**) Quantification of the flipping test based on a scoring system (n=7/group). (**G**) Quantification of the hind limb clasping test based on a scoring system (n=7/group). Data is depicted as mean ± standard deviation (SD), data points represent individual animals. For E, overall comparison was performed by two-way ANOVA with Tukey’s multiple comparison. Individual groups in E were compared using two-tailed student’s t-test. For F and G, one-way ANOVA with Tukey’s multiple comparison was performed. ns p > 0.05, **p < 0.01, ****p < 0.0001.

Given that motor impairment is a common symptom observed in FTLD-*GRN* patients and in DKO mice (*62–64*), our initial objective was to evaluate whether the administration of AAV expressing 8D3:PGRN could alleviate these phenotypes. Therefore, we conducted a longitudinal assessment of rotarod performance, commencing three days after AAV injection. While initially, all mice performed at a comparable level, WT animals increased time on the rotarod over 15 weeks regardless of treatment. In contrast, performance of both DKO and AAV-treated DKO mice decreased over time (Fig. 2E). However, AAV treatment significantly slowed the rate of decline in DKO mice, and in contrast to untreated DKO mice, AAV treated DKO mice never failed the test (rotarod time <3 s). To further assess motor function, we performed a flipping test at 15 weeks, evaluating righting ability. In this paradigm, saline-treated DKO mice could be flipped over, while most AAV-treated DKO mice could not be flipped or raised themselves immediately (Fig. 2F, Fig. S3A, Movie S1-4). We further conducted the hind limb clasping test (*42, 65*) and found that treated DKO mice showed significantly reduced hind limb clasping compared to untreated DKO mice, with some mice not exhibiting any clasping phenotype (score = 0) (Fig. 2G, Fig. S3B). In conclusion, AVV driven expression of 8D3:PGRN substantially lowered severe motor neuron deficits in the DKO mice.

### AAV(L):bPGRN reduces proteolysis defects, TDP-43 pathology and neurodegeneration

Based on the correction of functional motor deficits, we hypothesized AAV(L):bPGRN treatment may also correct the endolysosomal and autophagic protein degradation defects observed in the DKO mice (*42–44*). These mice present with an increase in ubiquitinated proteins in the brain (Fig. 3A,B), indicating defects in proteolysis by the ubiquitin/proteasome pathway. Moreover, the lysosomal sphingolipid-activator protein prosaposin, its functionally active cleavage product saposin D, and the autophagosome cargo protein p62 are all upregulated, suggesting endo-lysosomal and autophagy dysregulation (Fig. 3A,B) (*66, 67*). The accumulation of insoluble p62 and both LC3 I and its lipid modified form LC3 II, key players in the autophagy pathway (*67*), indicates the formation of pathological autophagic aggregates (Fig. 3A,B) (*68*). Immunofluorescence detection of p62 confirmed the presence of these aggregates (Fig. 3C,D). Strikingly, all these protein degradation-related phenotypes were significantly reduced in treated DKO mice compared to untreated animals (Fig. 3A-D).

**Fig. 3:**
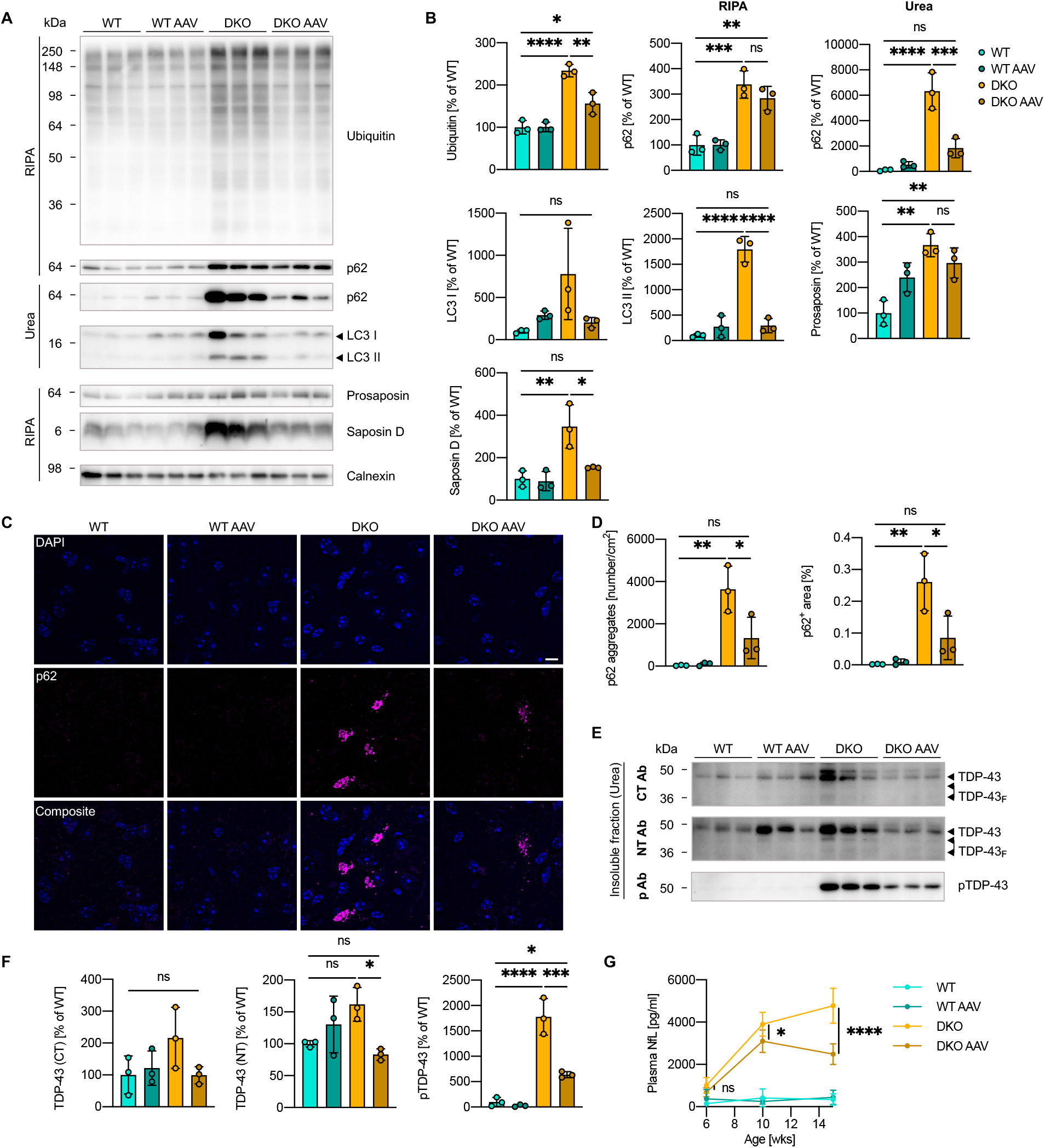
AAV-mediated delivery of 8D3:PGRN reduces protein degradation defects, TDP-43 pathology and NfL levels in DKO mice. (**A**) Western blot showing the protein levels of Ubiquitin, p62, prosaposin and saposin D in the RIPA fraction and the protein levels of p62 and LC3 I and II in the urea fraction in the brain of WT and DKO mice after treatment (n=3/group). (**B**) Quantification of the protein levels in A normalized to the mean of WT (n=3/group). (**C**) Representative immunofluorescence images of the thalamus of WT and DKO mice after treatment showing Map2^+^, Iba1^+^ and p62^+^ cells (scalebar = 10 µm). (**D**) Quantification of the number of p62 aggregates and p62^+^ area in whole brain hemispheres (n=3/group). (**E**) Western blot showing the protein levels of RIPA-insoluble TDP-43 in the urea soluble fraction of the brain of WT and DKO mice after treatment using antibodies directed against the C- and N-terminus as well as to the phosphorylation site Ser409/410 (n=3/group). (**F**) Quantification of the protein levels in E normalized to mean of WT (n=3/group). (**G**) Plasma NfL levels of mice before treatment (6 weeks) and 4 and 9 weeks after treatment (n=7/group). Data is depicted as mean ± SD, data points represent individual animals. For statistical analysis, one-way ANOVA with Tukey’s multiple comparison was performed. ns p > 0.05, *p < 0.05, **p < 0.01, ***p < 0.001, ****p < 0.0001.

DKO mice also exhibit TDP-43 pathology (*42–44*), a hallmark of FTLD-*GRN* that is observed in patients’ brains (*69, 70*), but is not robustly replicated in *Grn* KO mice (*11, 19*). This includes insoluble TDP-43 deposits, abnormal processing and phosphorylation of the protein (*42–44, 71*). Upon analyzing solubility, processing and phosphorylation of TDP-43 by sequential extraction and subsequent biochemical analysis, we only observed mild changes in protein levels and processing of TDP-43 in the high salt- and RIPA-soluble fraction (Fig. S4A-D). However, protein levels of insoluble TDP-43 (urea fraction) were consistently increased in DKO mice compared to WT mice, and this phenotype was completely reversed following AAV treatment (Fig. 3E,F). Additionally, we found a drastic increase of the pathological insoluble and phosphorylated TDP-43 (pSer409/410) in DKO mice compared to WT which was reduced by 70% in AAV(L):bPGRN-treated DKO mice (Fig. 3E,F). Because TDP-43 pathology is generally associated with neurodegeneration, we measured plasma neurofilament light chain (NfL) levels, a clinical marker for neurodegeneration (*7, 8*), before and during treatment. Initially, plasma NfL levels were comparable between WT and DKO mice. However, while NfL levels rose over time in DKO mice, they were significantly lower in treated DKO animals as early as 4 weeks after treatment and even decreased after 9 weeks (Fig. 3G). At 12 months of age, *Grn* KO mice only showed a mild increase in plasma NfL, but substantial increase of NfL in the CSF which was also rescued upon treatment with AAV(L):bPGRN (Fig. S4E).

Together, these findings demonstrate a robust reduction of TDP-43 pathology, associated with a rescue of the impaired protein degradation machinery, motor phenotypes, and reduced neurodegeneration following treatment with AAV(L):bPGRN.

### AAV(L):bPGRN reduces gliosis *in vivo*

Previous studies demonstrated that gliosis and neuroinflammation are hallmark pathologies not only in *Grn* KO (*12, 16, 19*) and the DKO mouse models (*42–44*), but also in FTLD-*GRN* patients (*5, 72*). To investigate if these pathological phenotypes could also be rescued by AAV(L):bPGRN treatment, we examined the transcriptional profile of *Grn* KO and DKO mice following AAV(L):bPGRN treatment. In 12-month-old *Grn* KO mice, bulk brain RNA sequencing analysis demonstrated mild transcriptional changes consistent with previous reports (*12, 15, 16, 19, 29*), which in turn were partially corrected with AAV(L):bPGRN (Fig. S5A). In 4-month-old DKO animals however, these changes were drastically exacerbated, consistent with previous reports (*42–44*). Projected into two dimensions via multi-dimensional scaling, samples from WT animals (with or without AAV treatment) clustered together, indicating globally similar gene expression profiles, but DKO samples segregated according to genotype and treatment (Fig 4A). AAV treatment of DKO animals shifted the corresponding samples toward those of WT animals, indicating partial correction of the transcriptional differences. We next completed gene-level differential expression analysis. We found that in DKO mice, 481 genes were up- and 304 genes downregulated compared to WT animals (Fig. S6A). Following AAV(L):bPGRN treatment, 51 genes were down- and 171 genes upregulated again in DKO mice (Fig. 4B). Pathway analysis revealed that the most impacted genes include established markers for disease-associated microglia (DAM) and lysosomal dysfunction (Fig. 4C) (*12, 73*). Interestingly, while the expression of many of these markers was fully corrected with AAV(L):bPGRN, some genes only showed partial response to treatment (ex:*Serpina3n*), suggesting *Tmem106b*-specific effects that are not responsive to PGRN replacement. Others, like *Ccl2*, showed an AAV response, regardless of genotype. As both DKO and treatment strongly modulated the abundance of known cell type markers such as *Gfap* (astrocytes) and *Itgax* (microglia) (Fig. 4B), we next determined changes in cell type composition (*74*) (Fig. S6B,C). Both, microglia and astrocyte markers were greatly increased in the DKO mice and showed a marked reduction with AAV treatment. In contrast, DKO brains had a reduction in neuronal cell type markers that are elevated with AAV treatment. Interestingly, different established oligodendrocyte markers displayed different gene expression changes across genotypes and treatments, perhaps reflecting differential responses in cellular subtypes. To confirm the significant reduction in gene expression associated with microglia and astrocyte activation following PGRN replacement, we further assessed the expression of Gfap, Iba1 and Cd68 by immunohistochemistry (Fig. 4D,E). In DKO mice, both the total number of microglia and Cd68^+^ reactive microglia, as well as astrocytes, were significantly elevated. Consistent with the RNA sequencing data, PGRN replacement resulted in a significant reduction of astrocytes. While the number of microglia was only slightly reduced, the proportion of Cd68^+^ microglia was substantially decreased (Fig. 4D,E). We further assessed the levels of Trem2, a microglial surface receptor that is critical for the transition to reactive microglia (*73, 75, 76*). The markedly elevated levels of Trem2 in plasma, liver and brain of DKO mice were all effectively decreased after PGRN replacement (Fig. 4F), consistent with a reduced microglial response to the pathological challenges after treatment-related amelioration. In *Grn* KO mice, comparable elevations of Cd68, Iba1 and Trem2 were also responsive to AAV(L):PGRN treatment, supporting the link between PGRN and glial function (Fig. S5B-D).

**Fig. 4:**
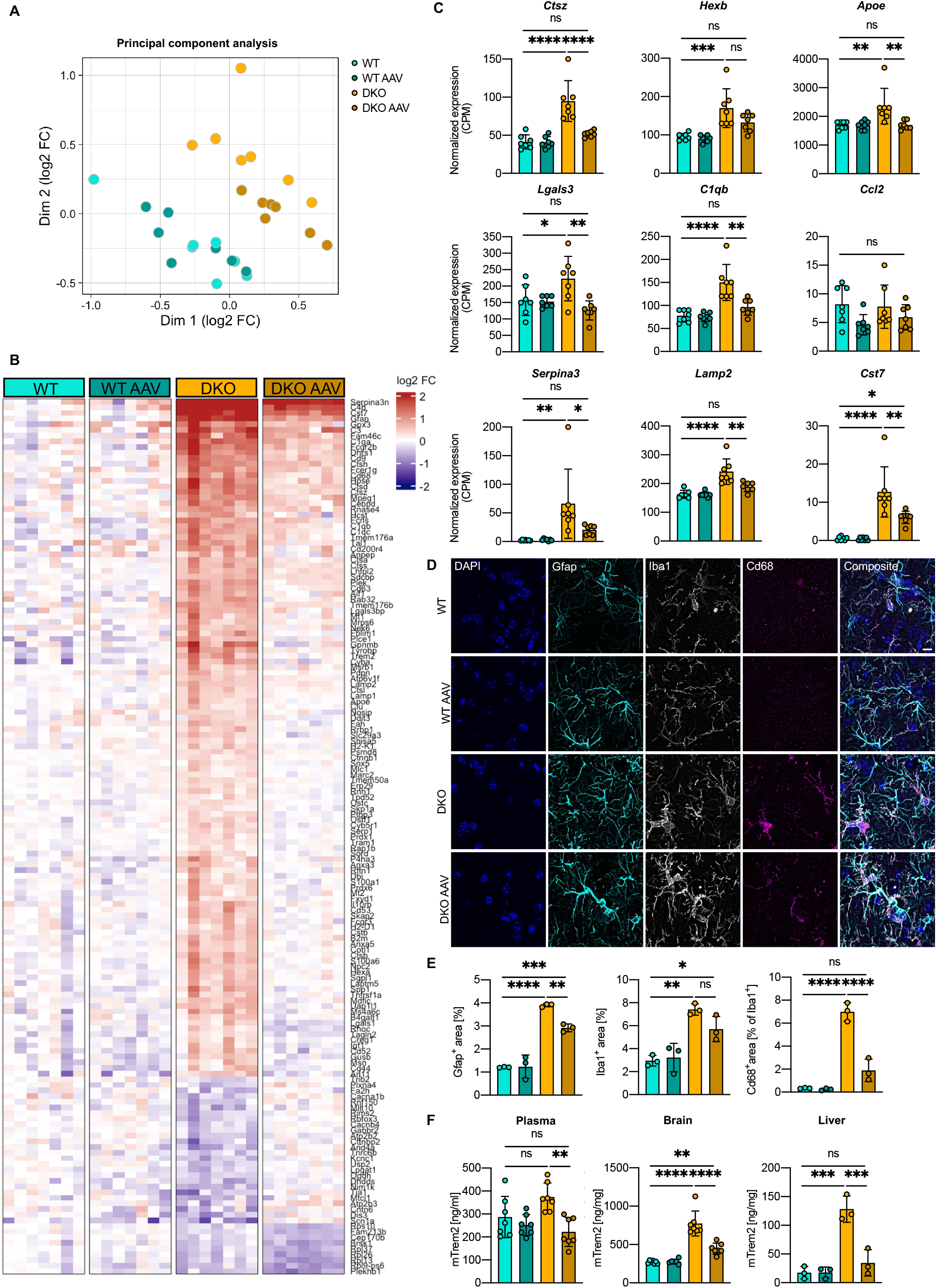
AAV-mediated delivery of 8D3:PGRN normalizes astrogliosis and microgliosis in DKO mice. (**A**) UMAP depiction of full transcript profile comparing untreated and AAV(L):bPGRN-treated WT and DKO mice (n=7/group). (**B**) Heatmap showing the top 222 differentially expressed genes in DKO mice and correction with AAV(L):bPGRN with a false discovery rate (FDR) < 5% and at least a 20% change comparing WT and DKO as well as DKO AAV and DKO. Plotted values are log2 transformed fold changes (FC) (n=7/group). (**C**) Transcriptional changes in genes linked to lysosomal function (n=7/group). (**D**) Representative immunofluorescence images of the thalamus of WT and DKO mice after treatment showing Iba1^+^, Gfap^+^ and Cd68^+^ cells (scalebar = 10 µm). (**E**) Quantification of the Gfap^+^, Iba1^+^ and Iba^+^/Cd68^+^ area in whole brain hemispheres (n=3/group). (**F**) mTrem2 levels in plasma (left, n=7/group), brain (middle, n=5-7/group) and liver (right, n=3/group) of WT and DKO mice after treatment at the end of the study as assessed by ELISA. C, E, F: Data is depicted as mean ± SD, data points represent individual animals. For statistical analysis, one-way ANOVA with Tukey’s multiple comparison was performed. ns p > 0.05, *p < 0.05, **p < 0.01, ***p < 0.001, ****p < 0.0001.

### AAV(L):bPGRN rescues lipid biomarkers of FTLD-*GRN*

Previously, *Grn* KO mice were found to exhibit robust alterations of lipid metabolism and lysosomal function (*29*). Here, we sought to determine if AAV(L):bPGRN can restore lipid changes. We first confirmed that 12-month-old *Grn* KO mice exhibit dysregulation of the phospholipid BMP and the glycosphingolipid glucosylsphingosine. Similar to recombinant PGRN replacement (*29*), AAV(L):bPGRN treatment restored these lipids back to baseline levels (Fig. S7A,B). Next, due to the established role of Tmem106b in the endolysosomal pathway (*46–50*), we hypothesized that the lipid profile of DKO mouse brain may be similarly or even more dysregulated. To assess this, we performed targeted lipidomic assessment of DKO and WT mouse brain with or without AAV(L):bPGRN treatment (Fig. 5). Unsupervised principal component analysis (PCA) of 203 lipids detected showed clear separation between WT and DKO mice along the principal component (PC) 1 axis. Lipid profiles of AAV(L):bPGRN-treated DKO mice shifted closer to WT conditions indicating a partial correction (Fig. 5A). Univariate analysis of lipid alterations in DKO mice brain relative to WT identified 27 significantly dysregulated lipids including several lipid classes such as gangliosides (GM1, GM2, GM3, GD3), glycerophospholipids (BMP, lysophosphatidylglycerol (LPG), phosphatidylinositol (PI)), sterols (cholesteryl ester, CE) and sphingolipids (S1P, Cer) (Fig. 5B,C). Of these, 19 lipids were responsive to AAV(L):bPGRN treatment with varying sensitivity (Fig. 5B-E). AAV(L):bPGRN treatment broadly increased BMP species of varying acyl-chain compositions back to WT levels (Fig. 5F). AAV(L):bPGRN treatment significantly lowered levels of galactosylsphingosine, which accumulates when galactosyl ceramidase (Galc) is impaired. The broad upregulation of gangliosides in *Grn* KO mice such as GM3, was also observed in DKO mice and further exacerbated by additional increase of GM2 and GM1 species and partially rescued upon treatment (Fig. 5G). The more robust ganglioside accumulating phenotype of DKO mice relative to *Grn* KO alone (Fig. S7) and only partial rescue upon AAV(L):bPGRN treatment suggests that TMEM106B may have a more dominant influence over lysosomal ganglioside catabolism. Also unique to DKO, there was an increase in myelin-associated lipids, CE and galactosylceramide (GalCer). Accumulation of these lipids potentially reflects the significantly altered myelin homeostasis in the DKO brain (Fig. 5H) (*42, 77, 78*). Taken together, these data highlight that DKO mice have severe lipid dysregulation that is largely corrected with PGRN replacement using AAV(L):bPGRN.

**Fig. 5:**
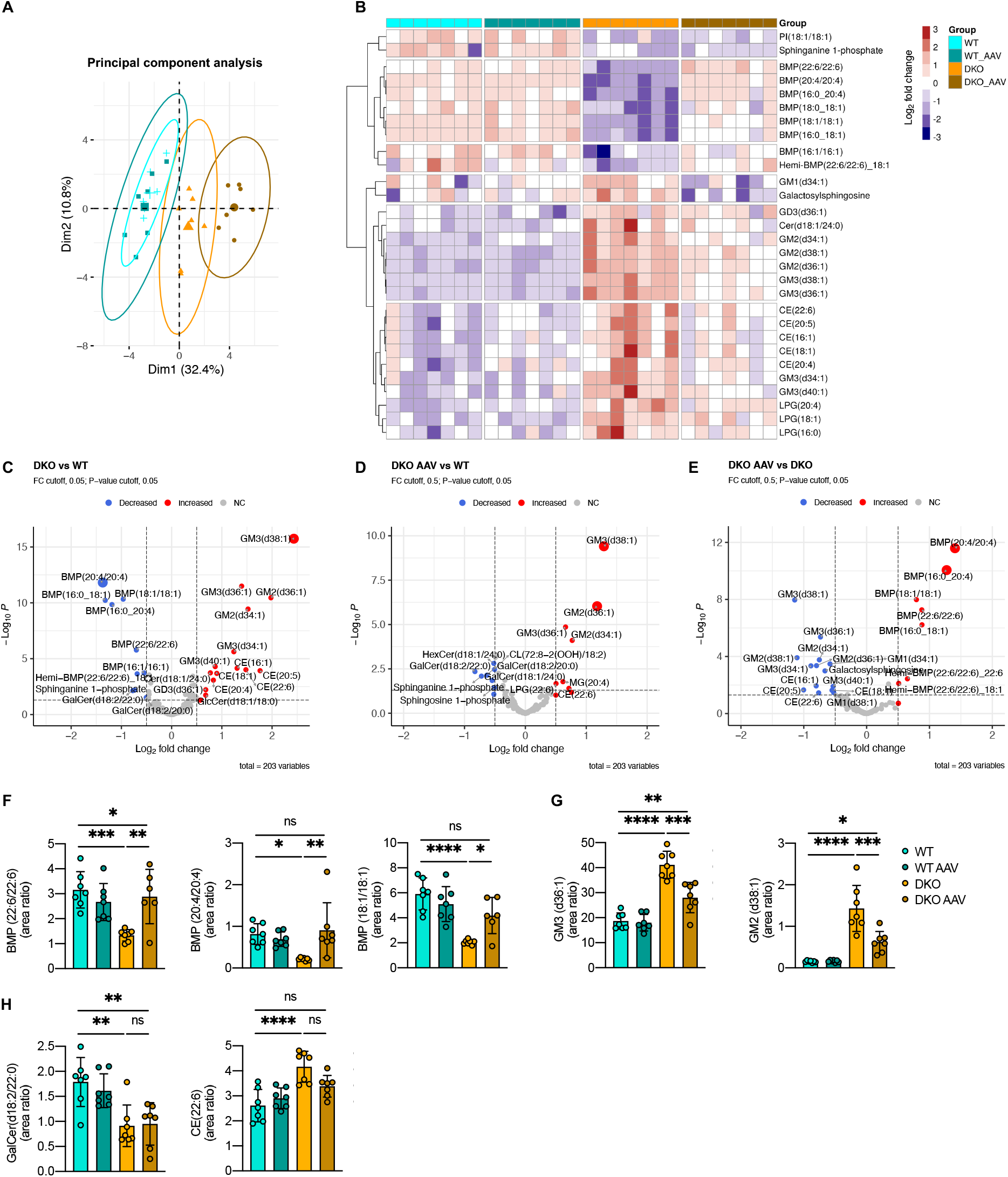
AAV-mediated delivery of 8D3:PGRN restores lipid profile of DKO mice. (**A**) Principal component analysis of global lipid changes comparing untreated and AAV(L):bPGRN-treated WT and DKO mice. (**B**) Heatmap depiction of differentially regulated lipids in DKO mice and correction with AAV(L):bPGRN treatment. Columns represent individual animals. Plotted values are log2 transformed. (**C-E**) Volcano plot depictions of major lipid changes that are genotype dependent (C) or treatment dependent comparing AAV-treated DKO mice (DKO AAV) to WT mice (D) or DKO mice (E). FDR cutoff: <5%, change cutoff: >20%. (**F-H**) Bar plots of treatment response of BMPs (F), gangliosides (G) and changes in myelin associated lipids (H). n=7/group. Data is depicted as mean ± SD ns p > 0.05, *p < 0.05, **p < 0.01, ***p < 0.001, ****p < 0.0001.

### PTV:PGRN rescues FTLD-like pathology in human neurons and microglia

To test whether our findings can be translated to human cells and to enable investigation of cell-autonomous versus non-autonomous effects, we extended our experiments to human induced pluripotent stem cells (hiPSC). First, we generated a DKO hiPSC line by knocking out *TMEM106B* in our recently described *GRN* KO cell line using an established CRISPR/Cas9 editing workflow (Fig. S8) (*79, 80*). To establish a hiPSC-based FTLD model, we co-cultured DKO hiPSC-derived microglia (iMG) with WT hiPSC-derived neurons (iN) (Fig. 6A). Crosstalk between these two cell types is expected as microglia express the highest PGRN levels of all brain cell types, whereas TDP-43 pathology is mostly observed in neurons (*71*). Upon co-culture of DKO iMG together with WT iN we observed a significant increase in TDP-43 processing and phosphorylation (Fig. 6B,C), resulting in pronounced disruption of the neuronal network as shown by a reduction in MAP2^+^ area (Fig. 6D,E). Lastly, we replicated our recently reported finding in *GRN* KO monocultured iMG showing dysfunctional lysosomes (*29, 32*). We found a strong elevation of the protein expression levels and catalytic activities of the lysosomal proteases cathepsin B, D and L (CTSB, CTSD, CTSL) in the co-cultures (Fig. 6F-H). As the *in vivo* mechanism of action of AAV(L):bPGRN cannot be replicated in a cell culture system, we applied PTV:PGRN (*29*) to the co-culture model to confirm therapeutic effects of PGRN replacement on TDP-43 pathology and neurodegeneration in a human system. Following a 5-day co-culture, we treated the cells with either PTV:PGRN, an inactive human IgG control (IgG) or PBS as control every second day until collection after 2 weeks (Fig. 6A). The cultures treated with PTV:PGRN showed partially normalized levels of PGRN, with fully reversed aberrant TDP-43 processing and disruption of the neuritic network (Fig. 6B-E). Moreover, the increased levels of pTDP-43 and lysosomal proteases as well as their increased activity were also reduced (Fig. 6B,C,F-H).

**Fig. 6:**
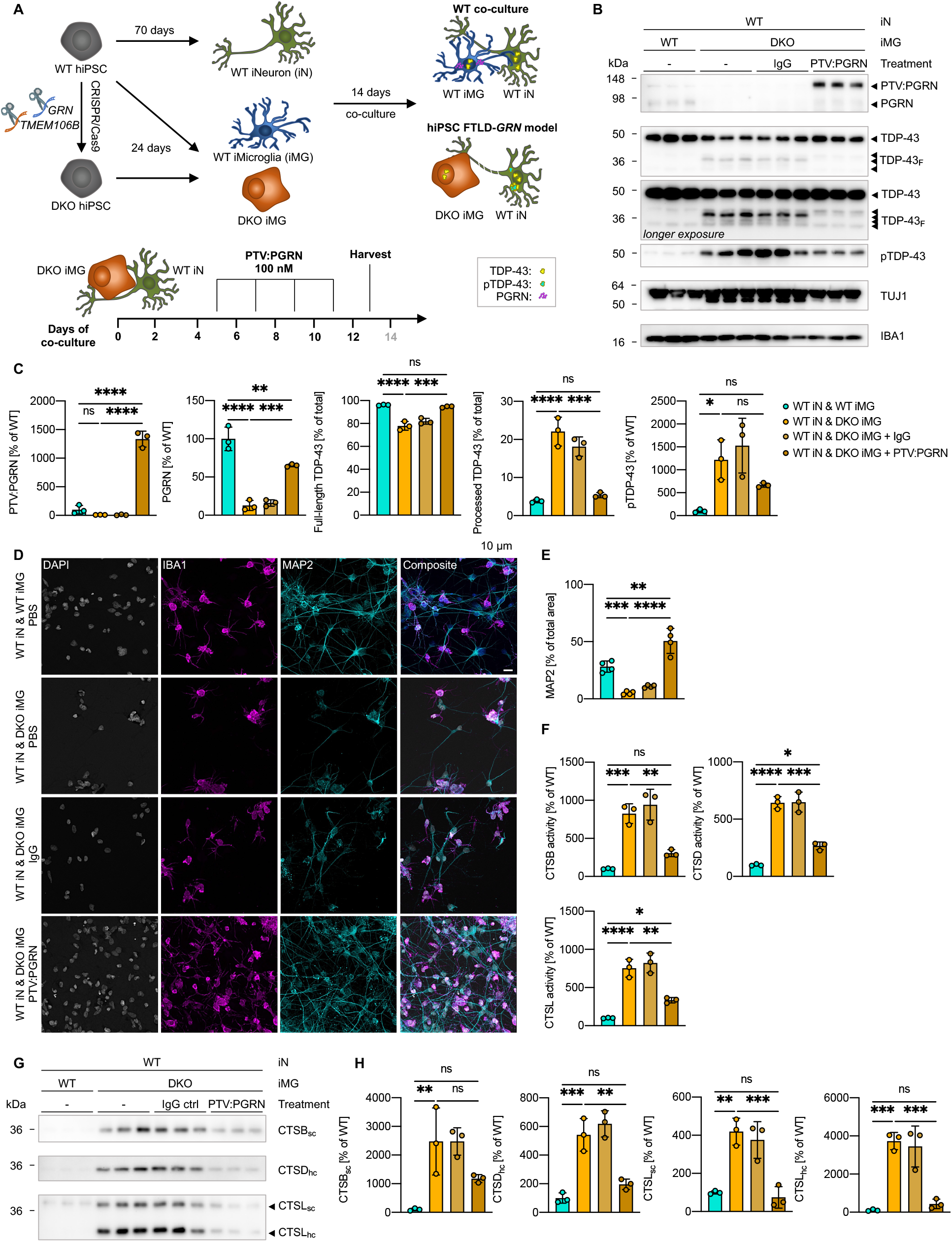
PTV:PGRN rescues FTLD-*GRN* core pathologies in hiPSC-derived DKO microglia in co-culture with neurons. (**A**) Graphical description of DKO hiPSC generation and differentiation, hiPSC-based co-culture FTLD-*GRN* model and treatment paradigm for PTV:PGRN treatment study. (**B**) Western blot showing the protein levels of PTV:PGRN and hPGRN, TDP-43 (NT Ab), pTDP-43, TUJ1 and IBA1 in untreated iMG / iN co-cultures as well as inactive control human IgG- and PTV:PGRN-treated cultures (n=3/biological replicates). (**C**) Quantification of the protein levels in B normalized to the mean of the WT iN / WT iMG co-culture (n=3/biological replicates). (**D**) Representative immunofluorescence images showing IBA1^+^ and MAP2^+^ cells (scalebar = 10 µm). (**E**) Quantification of the MAP2^+^ area (n=4/replicates). (**F**) *In vitro* activity of the lysosomal proteases cathepsin B (top left), cathepsin D (top right) and cathepsin L (bottom) normalized to the mean of the WT iN / WT iMG co-culture (n=3/biological replicates). (**G**) Western blot showing the protein levels of cathepsin B, D and L (n=3/biological replicates). (**H**) Quantification of the protein levels in G normalized to the WT iN / WT iMG co-culture (n=3/biological replicates). Data is depicted as mean ± SD, data points represent individual samples. For statistical analysis, one-way ANOVA with Tukey’s multiple comparison was performed. ns p > 0.05, *p < 0.05, **p < 0.01, ***p < 0.001, ****p < 0.0001.

Taken together, PGRN replacement in a human model successfully alleviates disease associated phenotypes and supports the applicability of our newly developed hiPSC-derived FTLD model for developing therapeutic approaches.

## DISCUSSION

Currently, there are several candidate therapeutic approaches for FTLD-*GRN* in clinical trials or preclinical development, all aiming to raise PGRN protein levels (reviewed in (*20, 55*)). These include an anti-sortilin antibody to increase circulating PGRN levels *via* inhibition of cellular uptake (*81–83*), AAV-based gene therapy approaches (*84–86*), a recombinant brain-penetrant PGRN biologic (*29*), as well as small molecules and transcriptional regulators (*87*). Here, we tested an innovative approach for a PGRN replacement therapy using a second-generation gene therapy approach in two different mouse models for FTLD-*GRN*, namely *Grn* KO and *GrnxTmem106b* DKO mice, that combines the dosing advantages of AAV approaches with the biodistribution improvements allowed by TfR-mediated transcytosis at the BBB and brain delivery of recombinant fusion proteins.

Despite several decades of research, mouse models reproducing TDP-43 pathology associated with FTLD-*GRN* without overexpression have been a major limitation in the field. Since *Grn* HET mice do not show any robust biochemical, pathological or behavioral phenotypes (*11, 40*), we took advantage of our recently developed model, which employs KO of *Grn* in combination with a KO of *Tmem106b* (*42*). This model was previously shown to reproduce abnormal phosphorylation and deposition of insoluble TDP-43, which are a central pathological hallmark of FTLD-*GRN*. This mouse model also exhibits profound motor phenotypes, in addition to exacerbating the phenotypes seen in *Grn* KO mice including deficits in protein degradation, lipid and transcriptome alterations, gliosis, and neurodegeneration (*42–44*).

In our study, a single injection paradigm resulted in successful AAV-mediated expression of a TfR-binding PGRN biologic (8D3:PGRN) in the periphery (*i.e.,* liver), which was robustly and sustainably delivered to the brain. This resulted in amelioration of the TDP-43 phenotypes within the brain. Furthermore, levels of endolysosomal lipid BMP, which was recently described as a pathway biomarker downregulated in *Grn* KO mice (*29, 33*), were fully restored. Levels of GM3 and GM2, which were upregulated in DKO mice were ameliorated as well, suggesting that the secondary storage of gangliosides observed in *Grn* KO mice is also responsive to PGRN replacement, despite the loss of *Tmem106b* (*29, 33*). Autophagic and lysosomal protein dysregulation was ameliorated as well. Transcriptomic and immunohistochemical analyses further indicated that microglia and astrocyte reactive states were significantly dampened. Altogether, these beneficial treatment effects were associated with a longitudinal decrease of plasma NfL, a biomarker for neurodegeneration which is commonly used in clinical practice (*8*). Lastly, the severe motor phenotypes in DKO mice, which started shortly after treatment, were substantially reduced. In contrast to other therapeutic strategies, which enhance circulating levels of PGRN by inhibiting cellular uptake (*e.g.*, the anti-sortilin antibody Latozinemab) (*83*), our approach increases both extracellular and intracellular PGRN. The latter is believed to be critical, as PGRN or its proteolytically generated granulins exert crucial functions within the endolysosomal pathway, which may be compromised by inhibition of cellular uptake of PGRN (*22, 88*).

We also verified the PGRN replacement strategy in a novel human iPSC-derived cellular model consisting of WT neurons co-cultured with *GRN/TMEM106B* DKO microglia. This model is based on the observation that *GRN* is predominantly expressed in microglia, whereas TDP-43 pathology primarily occurs in neurons (*71, 89, 90*). Microglial/neuronal crosstalk is in line with recent findings showing that complement factors secreted by *Grn* KO mouse microglia increase cell death and aberrant TDP-43 in mouse cortical neurons (*19*). Thus, we expected non-cell autonomous effects on neurons caused by loss of PGRN in microglia, which we indeed observed in our human co-culture system. Upon treatment with PTV:PGRN, disease-relevant abnormal posttranslational modification of TDP-43 and lysosomal dysfunction as well as neuronal loss were strongly ameliorated in the co-cultures, demonstrating applicability of this human model system to test efficacy of PGRN replacement.

Our toxicological evaluation of serum chemistry and hematology, as well as histopathological analysis did not show any significant adverse events and demonstrated that application of the brain-penetrant biologic AAV(L):bPGRN appears safe in mice at least for 31 weeks of treatment. This is of foremost importance, since mice intraventricularly injected with AAVs expressing PGRN showed neuroinflammation (*56*). Another caveat with protein replacement therapy is the impermeability of the BBB for AAVs or larger proteins, such as PGRN. Therefore, current therapeutic approaches investigated in the clinic rely on lumbar or cisterna magna CSF injection (*86*), which can cause injury to the medulla (*57*), and, despite direct delivery into the CNS, exhibit biodistribution limited to regions adjacent to CSF (*58*). This was also shown recently where direct CNS delivery of AAV1 and 9 expressing PGRN in a mouse model of FTLD-*GRN* resulted in amelioration of phenotypes in the spinal cord, but not in the brain (*60*).

Due to the severity of the DKO phenotypes, we employed a preventive subchronic treatment paradigm shortly before pathologies occur, while treating *Grn* KO animals in a chronic treatment paradigm. As genetic testing combined with measurements of plasma PGRN and NfL levels may allow for diagnosis of asymptomatic *GRN* mutation carriers before they convert to the clinical stage of the disease, prevention trials with PGRN replacement therapies may potentially be applied before clinical symptoms arise. Taken together, our findings strongly support the application of brain-penetrant PGRN as replacement therapy in FTLD-*GRN* patients. Furthermore, since our approach ameliorates lysosome-related deficits within the brain, similar brain penetrant therapeutics approaches may be developed for other lysosomal disorders with a neuronopathic component.

A limitation of our study is that by definition, the effect of PGRN replacement on TMEM106B fibrils (*51–53*) or of molecules aiming to increase PGRN levels, could not be addressed using this model due to the full loss of PGRN and TMEM106B. Another limitation of our and other models for TDP-43 pathology is the sparse TDP-43 mis-localization and nuclear clearance (*19, 42*). However, the biochemical TDP-43 phenotypes closely resemble those observed in patients (*71*).

## MATERIALS AND METHODS

### Animal experiments and mouse brain tissue

All animal experiments were performed either according to German animal welfare law and approved by the government of upper Bavaria (Vet_02-17-106) or by the Denali Therapeutics Institutional Animal Care and Use Committee. Mice were kept under standard housing conditions; standard pellet food and water was provided *ad libitum*. AAV was applied intravenously *via* tail vain injection (max. 200 µl). For tissue collection, mice were anesthetized with either Ketamine/Xylazine or tribromoethanol. Whole blood was either terminally collected via cardiac puncture or during the study via facial vein puncture and transferred into EDTA-coated tubes before centrifugation at 12,700 rpm for 7 min at 4°C. The plasma fraction was used for analysis. In cases where CSF was drawn, it was obtained by cisterna magna puncture and spun down at 12,700 rpm for 7 min at 4°C to minimize blood contamination. Mice were transcardiacally perfused with chilled PBS before organ collection. Tissues and fluids were stored at −80°C until further use.

### Generation and maintenance of WT and *GRNxTMEM106B* DKO iPSC lines

Experiments using hiPSC were performed following all relevant local guidelines and regulations. The iPSC line A18944 was purchased from Thermo Fisher (A18945). For the generation of *GRNxTMEM106B* DKO iPSC lines, our previously described *GRN* KO iPSC line was used (*32*). Cells were kept in Essential 8 Flex media (Thermo Fisher, A2858501) on vitronectin-coated (Thermo Fisher, A3349401) cell culture dishes at 37°C, 95% humidity and 5% CO_2_. Cells were detached as colonies at 70% confluency using PBS with 0.5 mM EDTA and passaged subsequently.

#### CRISPR/Cas9 genome editing

Human iPSC were edited as previously described (*80, 91*) with modifications. In brief, CRISPOR (http://crispor.tefor.net) (*92*) was used to design gRNAs and for the determination of putative off-target loci. To cover most of the coding region of the *TMEM106B* gene locus, exon 2 was targeted close to *de novo* STOP codons in alternate reading frames.

Cells were cultured as single cells on Geltrex (Thermo Fisher Scientific, A1110501) in StemFlex™ media (Thermo Fisher Scientific, A3349401) supplemented with ROCK inhibitor (Selleck-chem, S1049) for two days prior to electroporation. They were harvested using Accutase (Thermo Fisher Scientific, A1110501) and resuspended in 20 µl P3 nucleofector solution (Lonza) before addition to a pre-incubated ribonucleoprotein complex of gRNA (60 pmol) and Cas9 (30 pmol, Integrated DNA Technologies) protein. Using a Nucleofector (Lonza) with program CA-137, cells were electroporated and grown on Geltrex in StemFlex™ media with ROCK inhibitor until colonies appeared. Single colonies were picked and expanded upon Sanger sequencing verification of successful KO. Successful editing was further confirmed by Western blot. Quality controls were performed to confirm the absence of off-target effects (by PCR for the 10 most likely hits based on MIT and CFD scores on CRISPOR), the absence of loss of heterozygosity (nearby SNP sequencing), pluripotency of the novel cell line (immunofluorescence staining for OCT4, NANOG, SSEA4 and TRA160) and chromosomal integrity by molecular karyotyping (LIFE & BRAIN GmbH).

#### Differentiation of human iPSC-derived Microglia (iMG)

Cells were differentiated into iMG following the protocol initially presented by Abud et al. (*93*), with the modifications we recently published (*32*). On day 24 of differentiation, cells were used for co-cultures with iN.

#### Differentiation of human iPSC-derived Neurons (iN)

Cells were differentiated into cortical neurons (iN) according to our recently published protocol, following a dual-SMAD inhibition approach (*94*). On day 60 of differentiation, iN were used for co-cultures with iMG.

#### Co-culture of iMG and iN and PTV:PGRN treatment

For co-cultures, iMG and iN were cultured together at a 1:8 ratio. The required amount of iMG were collected and spun down as for passaging before resuspending in 50% of the required iN media, supplemented with double the concentration of the cytokines needed for iMG identity, namely hIL34 (Peprotech, 200-34), hM-CSF (Peprotech, 300-25) and hTGF-ß (Peprotech, 100-21). 50% of the media was removed from the iN and the fresh media containing the iMG added. From then on, 50% media without doubling the cytokine concentration was replaced every other day before cell processing for experiments on day 14 of co-culture.

To test the efficacy of PTV:PGRN in our model, cells were treated with either PBS or 100 nM inactive human IgG as a control, or 100 nM PTV:PGRN (*29*) starting on day 5 of co-culture every other day until collection.

### Design and generation of AAV(L):bPGRN

The mature protein expressed by AAV(L):bPGRN was designed by genetically fusing a mouse TfR binding scFv (8D3) to the N-terminus of full-length human PGRN. A human influenza hemagglutinin (HA) tag and 2x Gly4Ser linker were included between the scFv and PGRN to provide an additional handle for detection (HA tag) and to impart flexibility between the two proteins. Mature PGRN was used for the fusion protein, with N- and C-terminal breakpoints of T18 and L593, respectively; numbering as outlined in UniProt P28799. The fusion protein was cloned into a plasmid under an ApoE/AAT1 promoter, with a terminal WPRE to enhance expression, and flanking AAV2 ITRs. After generating the viral cis plasmid, recombinant AAV8 was generated and characterized by Signagen. The virus was titered by rtPCR quantification of genome copies. Endotoxin quantification was also performed and measured at less than 0.05 u/ml. Virus was delivered to animals as described above at a dose of 1.3×10^13^ vg/kg,

### Motor function assessment of DKO mice

All motor tests were performed blinded.

#### Rotarod test

Motor function was assessed using a rotarod (TSE Systems, 3376O4R), accelerating from 4-40 rpm within 300 s as previously described (*42*). Mice were trained twice within one week before longitudinal assessment twice per week. The average of three consecutive tests was calculated for each recording day.

#### Flipping test

Mice were recorded while being slightly poked to assess balance. Mice were scored based on showing no change in balance (0), trembling (1), flipping over and standing up within 3 seconds (2), or flipping over and not standing up within 3 seconds (3).

#### Hind limb clasping test

Mice were recorded while being lifted by their tail as previously described (*65*). They were scored based on no cramping of the hind limbs (0), cramping for less than 25% (1), more than 25% (2) or more than 50% of the recording time.

### Toxicity assessment

Assessment of toxicity was based on body weight, serum chemistry, hematology, necropsy and microscopic evaluation of select organs. Whole blood and serum were collected for complete blood count (CBC) and serum chemistry, respectively. Samples were evaluated at Quality Veterinary Laboratories (QVL). A complete necropsy was performed by qualified staff, and brain, heart, lung, liver, kidney, spleen, small and large intestine, prostate, testes, femur/tibia (including bone marrow) and sciatic nerve were collected into 10% buffered formalin. For tissue histopathology, tissues were processed to H&E-stained slides at Cureline Biopathology. Histology slides were reviewed by a board-certified veterinary pathologist, and instances of abnormal tissue observations were recorded.

### Gene profiling of total brain

Total RNA was extracted from frozen brain powder using the Qiagen RNeasy plus kit (Qiagen, 74034). Quality and quantity were assessed with a 4200 TapeStation System (Agilent). Next generation sequencing libraries were generated using the QuantSeq 3’ mRNA-seq Library Prep Kit FWD for Illumina sequencing with the UMI second strand synthesis module (Lexogen). Library concentration and quality were assessed with the 4200 TapeStation system (Agilent). The libraries were pooled in equimolar ratios for sequencing on an Illumina NovaSeq 6000 instrument. Raw sequencing data (FASTQ files) were processed with skewer (version 0.2.2) (*95*) to trim sequencing adapters. Unique molecular identifiers (UMIs) were extracted from each read with the umi2index tool (Lexogen). Quality control of the trimmed reads was performed using FastQC (version 0.11.5) (*96*). Reads were aligned to the mouse reference genome (version GRCm38_p6) with STAR (version 2.7.1a) (*97*), using an index created with the –sjdbOverhang=50 argument. Splice junctions from Gencode gene models (release M17) were provided via the –sjdbGTFfile argument. Alignments were obtained with the following parameters: –readFilesCommand zcat –outFilterType BySJout –outFilterMultimapNmax 20 –alignSJoverhangMin 8 –alignSJDBoverhangMin 1 – outFilterMismatchNmax 999 –outFilterMismatchNoverLmax 0.6 –alignIntronMin 20 – alignIntronMax 1000000 –alignMatesGapMax 1000000 –quantMode GeneCounts – outSAMunmapped Within –outSAMattributes NH HI AS nM MD XS –outSAMstrandField intronMotif –outSAMtype BAM SortedByCoordinate –outBAMcompression 6. Reads sharing the same UMI and genomic coordinate were deduplicated using the collapse_UMI_bam tool (Lexogen). Post-alignment quality control reports were generated using MultiQC (version 1.0) (*98*). The raw gene expression matrix was constructed from the ‘forward’ column of STAR’s ReadsPerGene.out.tab output files using R (version 4.3). Gene symbols and biotype information were extracted from the Gencode GTF file. For downstream analysis, library size factors were determined using TMM normalization (*99*) across all genes annotated as protein_coding with the edgeR R package (version 3.42.4) (*100*). Exploratory and differential expression analysis was performed with the edgeR and limma (version 3.56.2) (*101*) R packages. Multidimensional scaling was performed based on the top 500 most variable genes with the plotMDS function, after removing batch effects with removeBatchEffect function. For differential expression analysis, a linear model with fixed effects for experimental group, sex and batch covariates was fit to all protein-coding genes with the voomLmFit function, including sample weights. Contrasts of interest were extracted with the eBayes function with robust priors (e.g., by setting robust=TRUE). Genes with a false discovery rate (fdr) of less than 5% and a fold change of at least 20% were considered to display a significant genotype or treatment effect. Significant genotype x treatment interactions were called by applying a 15% fdr threshold. To visualize genes whose expression was perturbed in the double-knock out samples and also showed correction after AAV treatment, their normalized, batch-corrected expression was plotted with the ComplexHeatmap R package (version 2.16.0) (*102*). We found that in the DKO mice, *Grn* expression was still detected by our 3‘ mRNA sequencing approach, but protein PGRN was absent confirming successful knockout as previously described (*11, 15, 42*).

### Lipid analysis by liquid chromatography-mass spectrometry (LCMS)

Lipidomic analyses were largely performed as previously described (*29*). Approximately 20 mg tissue were homogenized for 30 sec at 25 Hz at 4°C in 400 µl of LC-MS grade methanol containing 2 µl internal standard mix solution. Homogenized samples were centrifuged at 21,000 g for 20 min at 4°C. Cleared methanolic supernatants were kept at −20°C for 1 h. Samples were centrifuged at 21,000 g for 10 min at 4°C and supernatants dried under a constant stream of N_2_. Dried samples were stored at −80°C until analysis. For analysis and separation of glucosylceramides and galactosylceramides, 50 µl of above extracts were dried under constant stream of N_2_ for 4 h and resuspended in 200 µl of 92.5/5/2.5 LCMS grade acetonitrile/isopropanol/water fortified with 5 mM ammonium formate and 0.5% formic acid.

Samples extracted above were analyzed using three different lipidomics assays, namely 1) lipidomics positive, 2) lipidomics negative and 3) glycosphingolipid positive panels. All of these assays were performed using the Agilent 1290 Infinity UPLC system coupled to Sciex 6500+ QTRAP mass spectrometer. All data acquisition was performed in multiple reaction monitoring mode (MRM) with the collision energy (CE) values reported in Table S2 and S3. Lipids were quantified using a mixture of non-endogenous internal standards as reported in Table S2 and S3. All quantification was performed using MultiQuant 3.02 software (Sciex). Chromatography and mass spectrometer settings used are described below:

Lipidomics positive: Mobile phase A consisted of 60:40 acetonitrile/water (v/v) with 10 mM ammonium formate + 0.1% formic acid; mobile phase B consisted of 90:10 isopropyl alcohol/acetonitrile (v/v) with 10 mM ammonium formate + 0.1% formic acid. Source settings for positive mode were as follows: curtain gas at 40 psi; collision gas was set at medium; ion spray voltage at 5500 V; temperature at 250°C; ion source Gas 1 at 55 psi; ion source Gas 2 at 60 psi; entrance potential at 10 V; and collision cell exit potential at 12.5 V.

Lipidomics negative: Mobile phase composition was same as in positive mode with the exception that 10 mM ammonium acetate and 0.1% acetic acid were used as fortificants. Source settings were as follows: curtain gas at 30 psi; collision gas was set at medium; ion spray voltage at −4500 V; temperature at 600°C; ion source Gas 1 at 55 psi; ion source Gas 2 at 60 psi; entrance potential at −10 V and collision cell exit potential at −15.0 V.

Lipidomics analysis in positive and negative ionization modes were initiated with 5 µl of sample injection using a BEH C18 1.7 μm, 2.1×100 mm column (Waters) as stationary phase with 0.25 mL/min flowrate and column temperature set at 55°C. The gradient was programmed as follows: 0.0-8.0 min from 45% B to 99% B, 8.0-9.0 min at 99% B, 9.0-9.1 min to 45% B, and 9.1-10.0 min at 45% B.

Glucosylsphingolipid panel in positive mode: Mobile phase A consisted of 92.5/5/2.5 ACN/IPA/H2O with 5 mM ammonium formate and 0.5% formic acid. Mobile phase B consisted of 92.5/5/2.5 H2O/IPA/ACN with 5 mM ammonium formate and 0.5% formic acid. For each analysis, 5 μL of sample was injected on a HALO HILIC 2.0 μm, 3.0 × 150 mm column (Advanced Materials Technology, PN 91813-701) using a flow rate of 0.48 mL/min at 45°C. Mobile phase A consisted of 92.5/5/2.5 ACN/IPA/H2O with 5 mM ammonium formate and 0.5% formic acid. Mobile phase B consisted of 92.5/5/2.5 H2O/IPA/ACN with 5 mM ammonium formate and 0.5% formic acid. The gradient was programmed as follows: 0.0–2 min at 100% B, 2.1 min at 95% B, 4.5 min at 85% B, hold to 6.0 min at 85% B, drop to 0% B at 6.1 min and hold to 7.0 min, ramp back to 100% at 7.1 min and hold to 8.5 min. Glucosylceramide and galactosylceramide were identified based on their retention times and MRM properties of commercially available reference standards (Avanti Polar Lipids, Birmingham, AL, USA). Table S4 shows specific analytes and internal standards used in this assay.

Integrated areas of endogenous lipids were divided by areas of spiked in stable isotope internal standards. Specific pairings of endogenous lipid to class level internal standards are described in Table S2-4. For all analysis, raw area ratio data was log2 transformed and filtered to retain variables that are present in at least 70% of samples. Missing values were imputed using the kNN method using the 5 nearest donor variables. Unwanted variations in the dataset were adjusted for using the RUV4 method. Genotype, treatment and genotype-:treatment interaction effects on lipids were assessed using a robust linear model with sex, batch, and RUV4 regression factors as covariates in the model. Top contrast results were visualized using heatmaps, boxplots or volcano plots. Bar plots of mean differences of selected analytes were created using GraphPad Prism. All analyses were performed using R statistical software (version 4.0.2; R Core Team 2020).

### Protein analysis and Western blotting

Mouse hemispheres were mechanically pulverized before sequential protein extraction as previously described (*11*).

Initially, brain samples were lysed in high salt lysis buffer (0.5 M NaCl, 10 mM Tris pH 7.7, 5 mM EDTA, 1mM DTT, 10% sucrose) and incubated on ice for 15 min followed by centrifugation. The supernatant was used as high salt fraction, the pellet was resuspended in RIPA lysis buffer (150 mM NaCl, 20 mM Tris pH 7.4, 2.5 mM EDTA, 0.1% (w/v) SDS, 1% (v/v) NP-40, 0.25% Na-deoxycholate) and incubated on ice for 15 min followed by centrifugation. The supernatant was used as RIPA fraction, the pellet was resuspended in urea lysis buffer (30 mM Tris pH 8.5, 7 M urea, 2 M thiourea, 4% CHAPS) and processed as before generating the urea fraction as supernatant after centrifugation. All lysis buffers were supplemented with PhosStop™ phosphatase inhibitor cocktail (Roche, 04906837001), protease inhibitor cocktail (Merck, P8340) and Benzonase^®^ (EMD Millipore, 70746-4).

Material from hiPSC was lysed in RIPA lysis buffer for 20 min on ice before centrifugation. Protein concentration was determined using BCA protein assay (Thermo Fisher Scientific, 23225). Equal amounts of protein were diluted in Laemmli buffer and separated by SDS-PAGE before transfer onto polyvinylidene difluoride or nitrocellulose membranes (both GE Healthcare Science). After primary antibody incubation (Progranulin (clone 8H10, (*11*)), Calnexin (Stressgen, SPA-860), Ubiquitin (Cellsignaling, 3936), p62 (MBL, PM045), LC3 (Novusbio, NB100-2220), Saposin D ((*103*)), TDP-43 NT (Proteintech, 10782-2-AP), TDP-43 CT (Proteintech, 12892-1-AP), pTDP-43 (Proteintech, 80007-1-RR), b-Actin (Sigma-Aldrich, A5316), Progranulin (Thermo Fisher Scientific, 40-3400), TUJ1 (Biolegend, 801201), IBA1 (Gentex, GTX100042), Cathepsin B (Santa Cruz, Sc-13985), Cathepsin D (R&D Systems, AF1029), Cathepsin L (R&D Systems, AF1515), TMEM106B (clone 344, (*104*)), GAPDH (Invitrogen, AM4300)) followed by incubation with HRP-conjugated secondary antibodies, detection using ECL Plus substrate (Thermo Fisher Scientific) and imaged using an Amersham ImageQuant 800 system (Cytiva). Images were quantified using the MultiGaugeV3.0 software (Fujifilm Life Science).

### Lysosomal protease activity assays

Cells were lysed under mild conditions (50 mM Na-citrate pH 5.0, 0.8% (v/v) NP-40) for 15 min on ice before centrifugation and lysosomal protease activities performed as previously described (*12*). Activity assays for cathepsin B (Abnova, KA0766), cathepsin D (Abnova, KA0767) and cathepsin L (Abnova, KA0770) were performed according to the manufacturer’s instructions using samples with normalized protein concentrations. Fluorescence generated by cleavage of the quenched substrate was measured using a Fluoroskan Ascent FL plate reader.

### ELISA-based quantification of mTrem2

For the analysis of total cellular and soluble Trem2, samples were homogenized in either high salt lysis buffer or CST lysis buffer (Cell Signaling Technologies (CST), 9803) with protease and phosphatase inhibitors. Trem2 measurements were made using the Mesoscale Discovery (MSD) platform as previously described (*29, 105*) using a biotinylated anti-mTrem2 polyclonal antibody (R&D Systems BAF1729) as capture antibody and a sulfo-tagged detection antibody (sheep anti-mouse Trem2, AF1729 R&D Systems). MSD plates were developed using 2x MSD read buffer T, followed by detection using an MSD Meso Sector S600 reader. MSD values were converted to absolute quantities of Trem2 by interpolating from a 4-parameter logistic curve fit to the mouse Trem2-His protein standard using Graphpad Prism software. For the same samples BCA protein analysis was performed and Trem2 levels were normalized using the protein concentrations.

### ELISA-based quantification of mPGRN and hPGRN

For the analysis of total cellular and soluble PGRN in DKO mice, samples were homogenized in high salt lysis buffer with protease and phosphatase inhibitors. PGRN measurements were made using the Mesoscale Discovery (MSD) platform as previously described (*106*) using either a biotinylated anti-hPGRN antibody (R&D Systems, BAF2420) or anti-mPGRN antibody (R&D Systems, BAF2557) at 0.2 µg/mL as capture antibody and for detection rabbit anti-hPGRN (Thermo Fisher Scientific, 40-3400) or a rat anti-mPGRN (R&D Systems, MAB25571) antibody at 0.25 µg/mL followed by sulfo-tagged secondary antibodies (anti-rabbit and anti-rat respectively). MSD plates were developed using 2x MSD read buffer T, followed by detection using an MSD Meso Sector S600 reader. MSD values were converted to absolute quantities of PGRN by interpolating from a 6-parameter logistic curve fit to the recombinant protein standard (Workbench v4 software (MSD)).

Human PGRN levels in *Grn* KO mice were measured using the DuoSet ELISA kit (R&D DY2420) as previously described (*29*). Concentrations were calculated by interpolating values against an 8 point standard curve using Graphpad Prism.

### Neurofilament light-chain analysis

Plasma and CSF neurofilament light chain (NfL) levels were analyzed as previously described (*29*) using the Quanterix Simoa Neurofilament Light Advantage kit. Plasma was diluted 20x in sample diluent and CSF was diluted 100x in sample diluent. Sample NfL concentrations were measured using the NfL analysis protocol on the Quanterix SR-X instrument and interpolated against a calibration curve provided with the Quanterix assay kit.

### Immunofluorescence labeling of mouse brain tissue

For the analysis of DKO mouse brains, tissue was postfixed with 4% paraformaldehyde (PFA) for 24 h directly after tissue extraction, cryoprotected in 30% sucrose for 3 nights and subsequently embedded in Tissue-Tek OCT cryo-embedding matrix prior to freezing the tissue on dry ice. Brain samples were cut into 16 µm sections (CryoStar NX70 HOMVP, Thermo Fisher Scientific) and stored at −80°C. Immunofluorescence staining was performed on sections that were adjusted to RT for 30 min and washed with PBS to remove excess embedding matrix. Sections were transferred to citrate buffer (0.01 M, pH 6.0) for antigen retrieval. Tissue permeabilization was performed by incubating sections with 0.5% Triton-X-100 in PBS for 10 min. Blocking was performed using a mixed blocking solution (2.5% bovine serum albumin, 2.5% fish gelatin, 2.5% fetal calf serum in PBS). Primary antibodies (Map2 (1:500, Novusbio, NB300-213), Iba1 (1:400, Synaptic Systems, 234004), Gfap (1:500, Thermo Fisher Scientific, 13-0300) Cd68 (1:500, abcam, ab125212), p62 (1:200, MBL, PM045)) were diluted in antibody diluent (0.625% bovine serum albumin, 0.625% fish gelatin, 0.625% fetal calf serum in PBS) and incubated for 24 h at 4°C, followed by incubation with fluorophore-coupled secondary antibodies (Thermo Fisher Scientific), for 1 h at RT. Afterwards, sections were incubated with DAPI (1:1000, Thermo Fisher Scientific, 62248) for 10 min and mounted with ProLong™ Gold Antifade mounting solution (Invitrogen, P36930).

For the analysis of *Grn* KO mouse brains, samples were processed as previously described (*29*), tissue was postfixed with 4% PFA for 24 h directly after tissue perfusion and extraction, cryoprotected in 30% sucrose overnight before section at 40 µm thickness using a freezing microtome. Immunofluorescence staining was performed on free floating sections. Blocking and antibody washes were performed using a mixed blocking solution (1% BSA + 5% Normal Donkey Serum + 0.3% Triton-X-100 in PBS). Primary antibodies (Cd68 (1:500, BioRad MCA1957), Iba1 (1:400, Novus NB100-1028), and Gfap (1:500, Novus NBP1-05198) were used and incubated for 24 h at 4°C, followed by incubation with fluorophore-coupled secondary antibodies (Thermo Fisher Scientific), for 2-4 h at RT. Afterwards, sections were incubated with DAPI (1:5000, Thermo Fisher Scientific, 62248) for 10 min and mounted with ProLong™ Gold Antifade mounting solution (Invitrogen, P36930).

### Immunofluorescence labeling of hiPSC cultures

Cells were fixed by replacing the culture media with 4% PFA (Thermo Fisher Scientific, J19943.K2) and incubating at 37°C, 95% humidity and 5% CO_2_ for 15 min, before storage at 4°C in PBS until further processing as previously described (*107*). After two initial PBS washes, cells were permeabilized using 0.1% Triton-X-100 in PBS, before blocking in SuperBlock™ blocking buffer (Thermo Fisher Scientific, 37515). Primary antibodies (MAP2 (1:1000, abcam, ab92434), IBA1 (1:400, Invitrogen, PA5-27436)) were diluted in SuperBlock™ and samples incubated at 4°C for 24 h followed by incubation with fluorophore-coupled secondary antibodies (Thermo Fisher Scientific) for 90 min at RT. Cells were incubated with DAPI for 10 min and mounted with ProLong™ Gold Antifade mounting solution.

### Image acquisition

For quantification, full brain hemispheres were imaged at 20x using a Zeiss Axio Scan.Z1 digital slide scanner or a Leica DMI-8 epifluorescence microscope with a DFC9000GT camera and a 20x objective (HC PL FL L 20x/0.40 CORR PH1). Image mosaics were merged using the Leica LAS X software suite. Analysis was completed using Zeiss Zen Blue 3.2 software or ImageJ. Image pre-processing to eliminate illumination gradient aberrations using a difference of gaussian filters with the CLIJ2 plugin (*108*) was performed. Regional ROIs were drawn, and a rolling ball thresholding approach was used to determine the area of each marker relative to total regional area. 1-3 sections were analyzed per brain and average percent coverage values were calculated across images. Representative images were acquired on a Zeiss spinning disc microscope using a 40x/1.3 n.A. oil immersion objective or on a Zeiss LSM800 Axio Observer 7 confocal microscope with a 63x (63x/1.4 Oil) or a 40x (40x/1.3 Oil) objective. Post-processing was performed in ImageJ.

### Statistical analysis

All non-sequencing or lipidomics data was analyzed using either a Student’s t-test, one-way analysis of variance 1555 (ANOVA), or two-way ANOVA with multiple comparisons when indicated. Data was plotted and statistical tests were performed using GraphPad Prism. All studies and sample analyses were performed blinded and in randomized order. Studies with multiple assays were designed to be powered for the most variable analyte based on previous data. Exclusion of mice only occurred if confounding health concerns were identified during the study. Data was only excluded in circumstances when extreme outliers were identified (ROUT, Q = 0.5%). All heatmaps were generated in R using the Complex Heatmap package. Data represented as mean ± SEM or mean ± SD for *in vitro* and *in vivo* graphs. N is defined as the number of independent experiments or biological replicates in an *in vitro* study and as the number of mice in an *in vivo* study.

## Acknowledgements

We would like to thank Dr. Masugi Nishihara (Department of Veterinary Physiology, University of Tokyo) for providing the *Grn* KO founder mouse strain used at DZNE Munich and Dr. Benedikt Wefers (Taconic Biosciences) and Dr. Wolfgang Wurst (Helmholtz Center Munich) for the generation of the *Grn/Tmem106b* DKO mouse strain. We want to thank Dr. Astrid Feiten for helping with analysis of behavior experiments, Dennis Crusius for help with iPSC experiments, Ramona Rodde for mouse genotyping, Claudia Thiel for help in rotarod assessment (all Ludwig Maximilian University of Munich) and all employees of the animal facility of the DZNE Munich for animal maintenance. We thank Martin Larhammar, Mollie Jacobs, and Zach Sweeney for contributions in the early stage of the project, as well as Connie Ha, Timothy K. Earr, Hoang Nguyen, and Roni Chau for technical help. We also thank Rene Meisner for the toxicology analysis, Karen Lai, Richard Tsai and our Takeda colleagues for critical reading of the manuscript.

## Author Contributions

Conceptualization: MR, MJS, MSK, SLD, JWL, AC, GDP, CH

Methodology: MR, MJS, IP, GW, SR, SSD, GL, LSc, LSp, SB, TL, MS, MSK, JHS, TS, SLD, JWL, DP, AC, GDP, CH

Investigation: MR, MJS, BP, CS, IP, GW, SR, SSD, GL, LSc, LSp, SB, BN, KB, JHS, TS

Visualization: MR, MJS, IP, JHS, TS

Funding acquisition: DP, CH

Project administration: DE, MS, JWL, DP, AC, GDP, CH

Supervision: MR, MJS, GW, SR, JHS, TS, LSc, LSp, SB, DP, AC

Writing – original draft: MR, MJS, GDP, CH

Writing – review & editing: MR, MJS, BP, GW, IP, LSp, TL, MS, JHS, TS, MSK, SLD, JWL, DP, AC, GDP, CH

## Data and materials availability

The bulk RNA sequencing data and lipidomics data will be made available in an online repository.

## Supplementary Materials

**Fig. S1:**
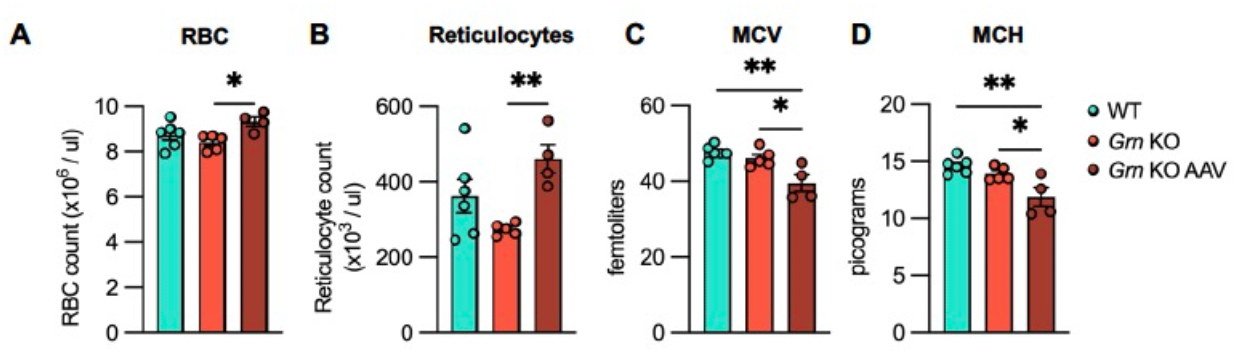
AAV-expressed 8D3:PGRN has minimal effects on red blood cells in *Grn* KO mice. Quantification of blood cell measurements including number of red blood cells (RBC) (**A**), number of reticulocytes (**B**), mean corpuscular volume (MCV) (**C**) and mean corpuscular hemoglobin (MCH) (**D**) in 12-month-old *Grn* KO mice following 31 weeks of AAV(L):bPGRN treatment. Data is depicted as mean ± SEM, data points represent individual samples. For statistical analysis, one-way ANOVA with Tukey’s multiple comparison was performed. *p < 0.05, **p < 0.01.

**Fig. S2:**
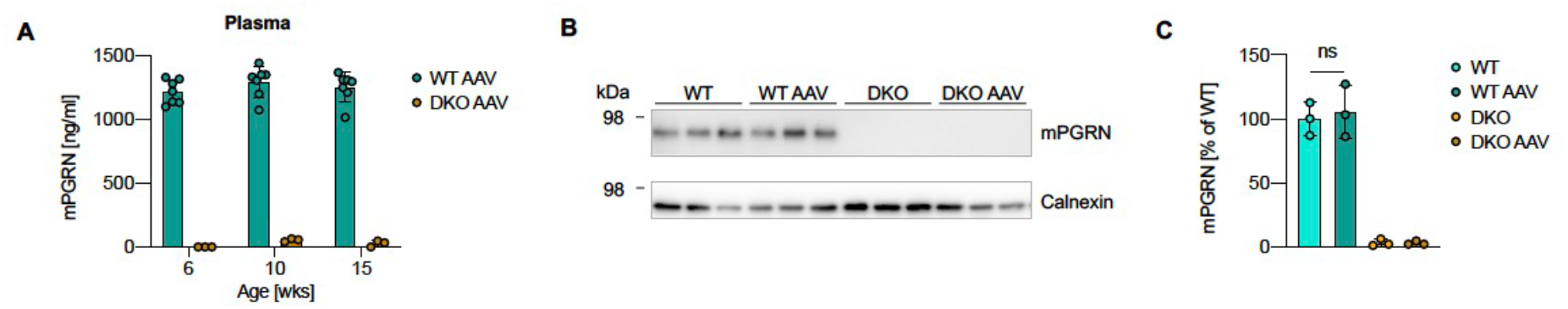
AAV-mediated delivery of 8D3:PGRN does not alter endogenous PGRN levels in WT mice. (**A**) mPGRN levels in plasma following AAV(L):bPGRN treatment in WT and DKO mice as assessed by ELISA (n=3-7/group). (**B**) mPGRN levels in brain following treatment as assessed by Western blot (n=3/group). (**C**) Quantification of mPGRN levels in B normalized to WT (n=3/group). Data is depicted as mean ± SD, data points represent individual animals. For statistical analysis, one-way ANOVA with Tukey’s multiple comparison was performed. ns p > 0.05.

**Fig. S3:**
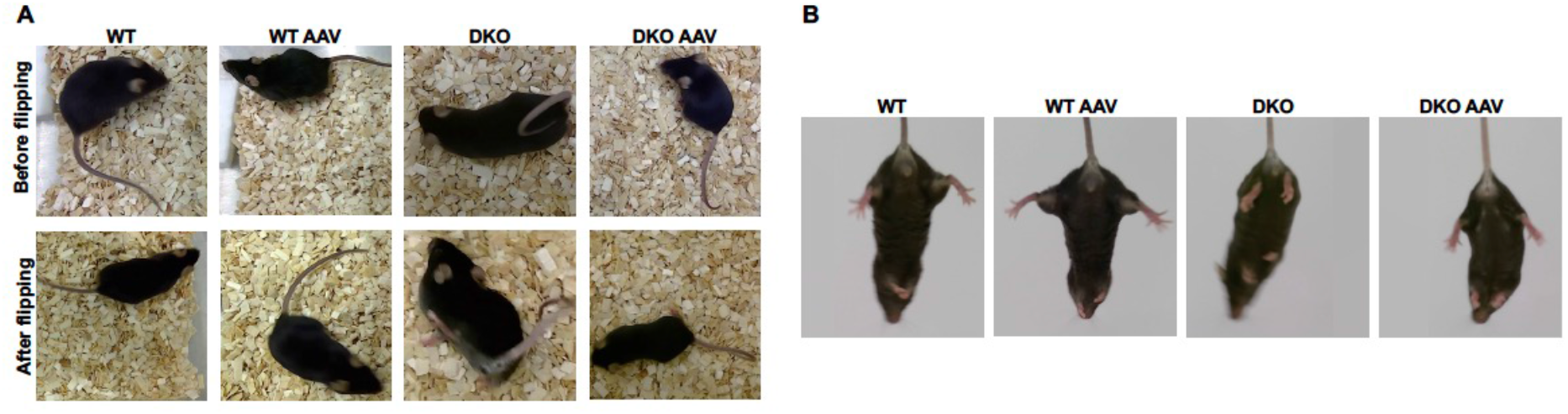
AAV-expressed 8D3:PGRN ameliorates behavioral abnormalities in DKO mice. (**A**) Representative images of 15-week-old mice 9 weeks after AAV(L):bPGRN treatment before and after flipping. (**B**) Representative images of hind limb clasping test showing 15-week-old mice 9 weeks after treatment.

**Fig. S4:**
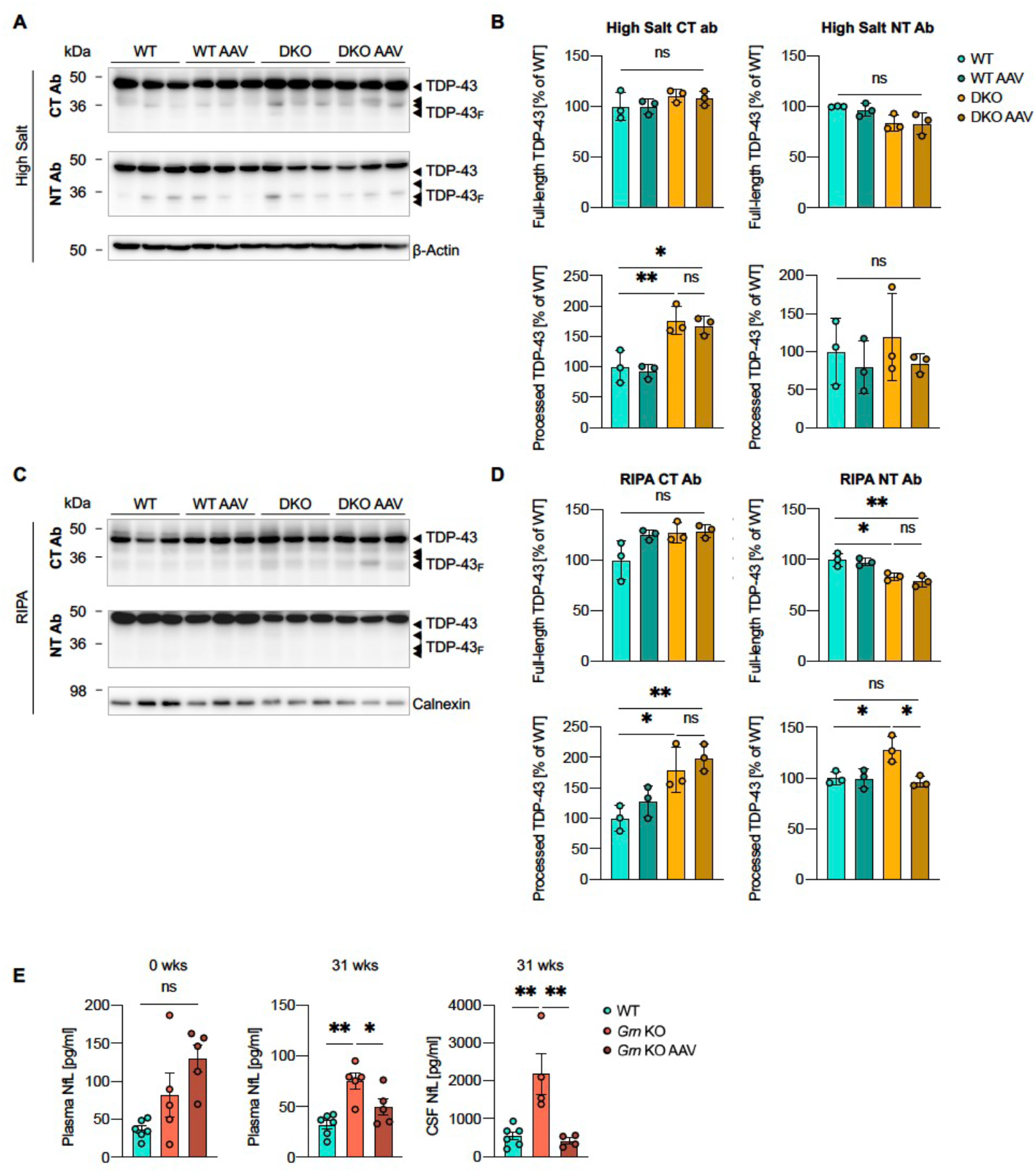
AAV-mediated delivery of 8D3:PGRN reduces TDP-43 pathology in DKO and NfL in *Grn* KO mice. (**A,C**) Western blot showing the protein levels of TDP-43 in the high salt (A) and RIPA (B) fractions of brains of untreated and AAV(L):bPGRN-treated WT and DKO mice by using antibodies either directed against the C-terminus or the N-terminus (n=3/group). (**B,D**) Quantification of the protein levels in A (B) and C (D) normalized to WT (n=3/group). (**E**) NfL levels in plasma and CSF of *Grn* KO animals following AAV(L):bPGRN treatment. n=5-6/group. Data is depicted as mean ± SD (B,D) or mean ± SEM (E), data points represent individual animals. For statistical analysis, one-way ANOVA with Tukey’s multiple comparison was performed. ns p > 0.05, *p < 0.05, **p < 0.01.

**Fig. S5:**
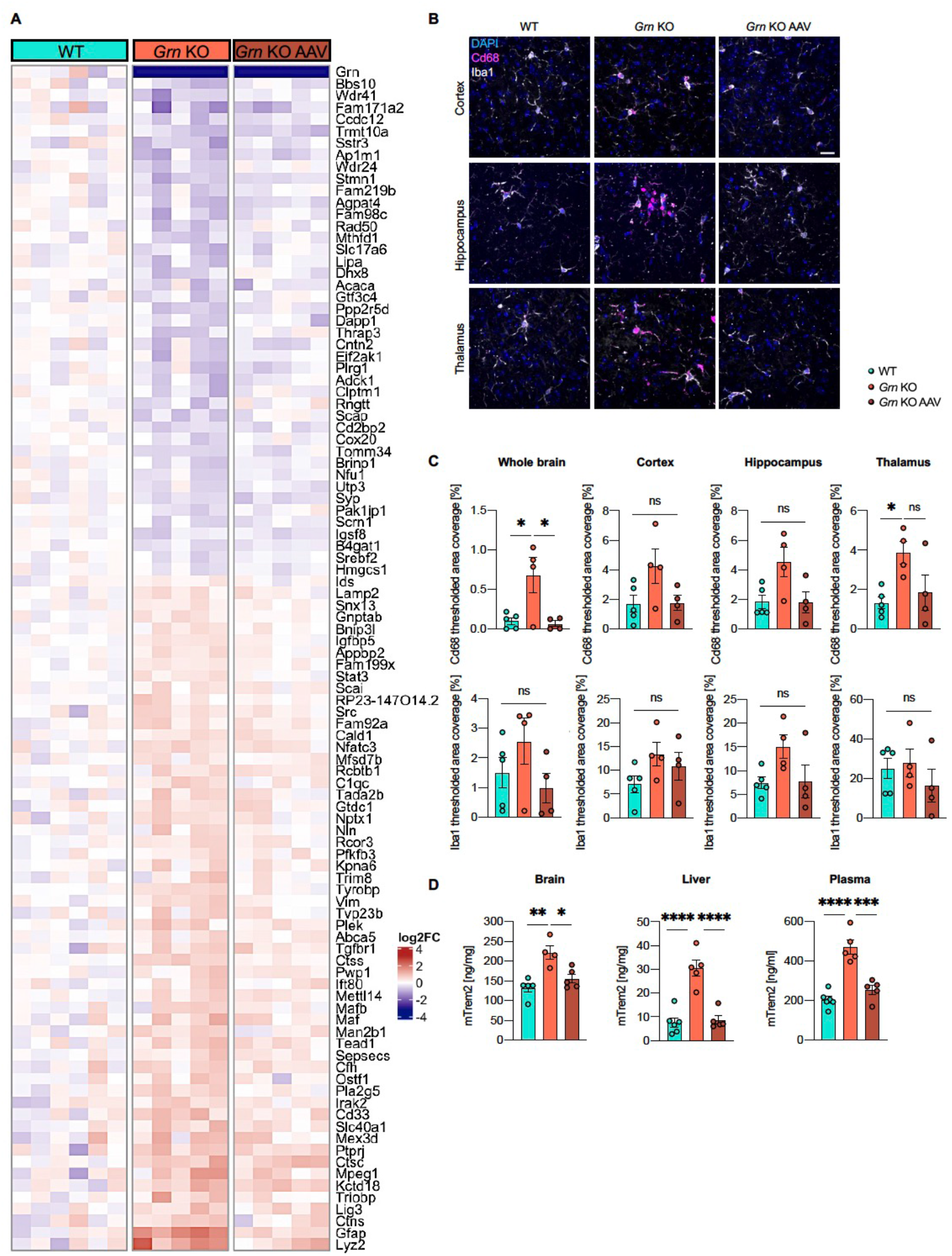
AAV-mediated delivery of 8D3:PGRN normalizes astrogliosis and microgliosis in *Grn* KO mice. (**A**) Heatmap depiction of transcriptional changes in 12-month-old *Grn* KO mice following 31 weeks of AAV(L):bPGRN treatment. Columns represent individual animals. Plotted values are log2 transformed. (**B**) Representative immunofluorescence images of the thalamus showing Iba1^+^ and Cd68^+^ cells (scalebar = 20 µm). (**C**) Regional quantification of Cd68 and Iba1 in whole hemispheres (left), cortex (middle left), hippocampus (middle right) and thalamus (right). (**D**) ELISA-quantified mTrem2 levels in brain (left), liver (middle) and plasma (right) in 12-month-old *Grn* KO mice following 31 weeks of AAV(L):bPGRN treatment. n=5-6/group. Data is depicted as mean ± SEM ns p > 0.05, *p < 0.05, **p < 0.01, ***p < 0.001, ****p < 0.0001.

**Fig. S6:**
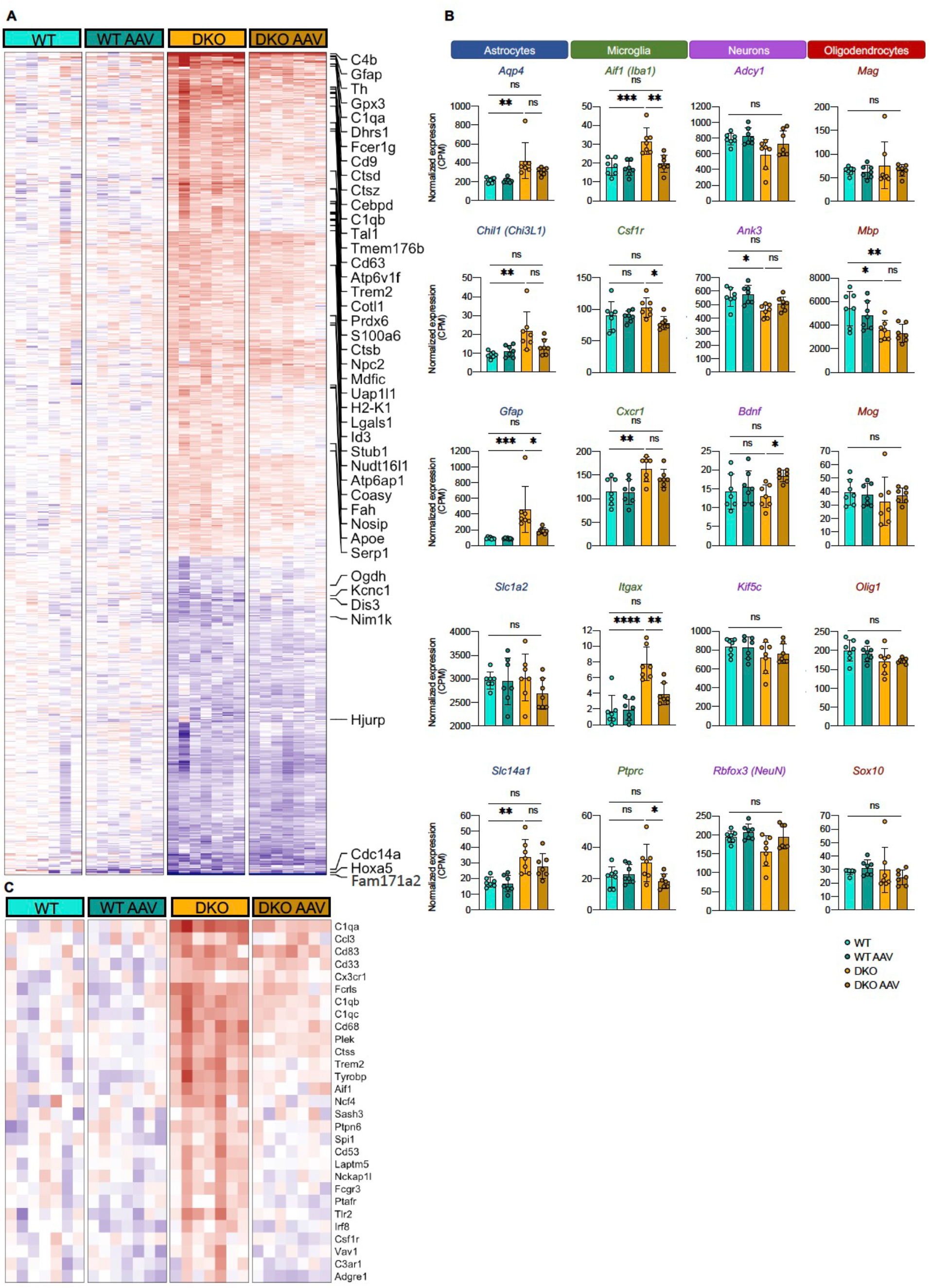
AAV-mediated delivery of 8D3:PGRN partially restores genetic cell type composition within the brain. (**A**) Heatmap depiction of the top 785 differentially expressed genes in DKO mice and correction with AAV(L):bPGRN with a false discovery rate (FDR) < 5% and at least a 20% change comparing WT and DKO. Plotted values are log2 transformed. (**B**) Expression of astrocyte (blue), microglia (green), neurons (purple) and oligodendrocyte (red) markers in the treated and untreated WT and DKO mice. (**C**) Heatmap depiction of the top 29 differentially expressed microglia-specific genes in DKO mice and correction with AAV(L):bPGRN with a false discovery rate (FDR) < 5% and at least a 20% change comparing WT and DKO. Plotted values are log2 transformed. n=7/group. Data is depicted as mean ± SD. ns p > 0.05, *p < 0.05, **p < 0.01, ***p < 0.001, ****p < 0.0001.

**Fig. S7:**
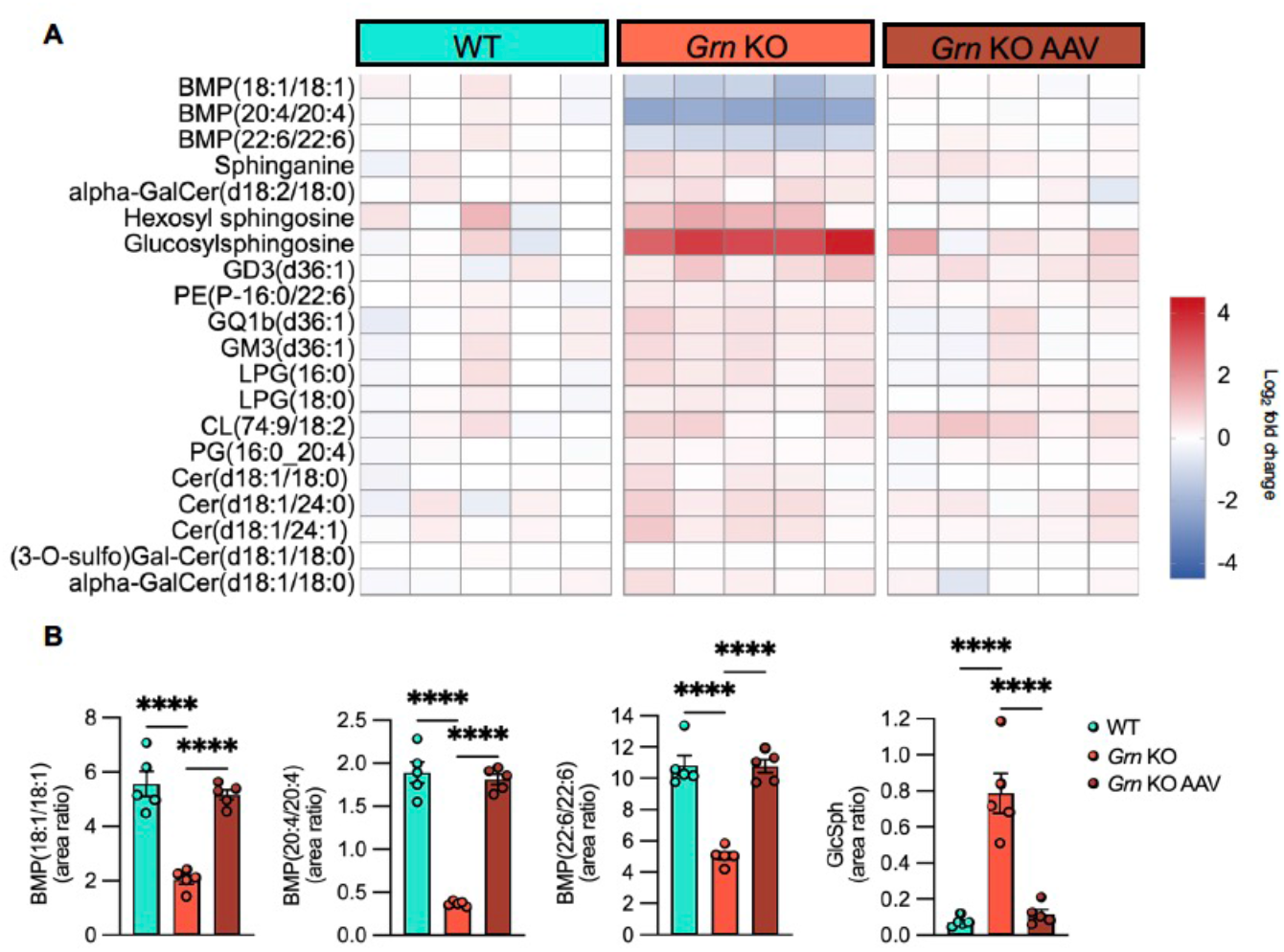
AAV-mediated delivery of 8D3:PGRN restores lipid profile of *Grn* KO mice. (**A**) Heatmap depiction of differentially regulated lipids in *Grn* KO mice and correction with AAV(L):bPGRN treatment. Columns represent individual animals. Plotted values are log2 transformed. FDR cutoff: <5%, change cutoff: >20%. (**B**) Bar plots of treatment response of BMPs and gangliosides. n=5/group. Data is depicted as mean ± SEM. ****p < 0.0001.

**Fig. S8:**
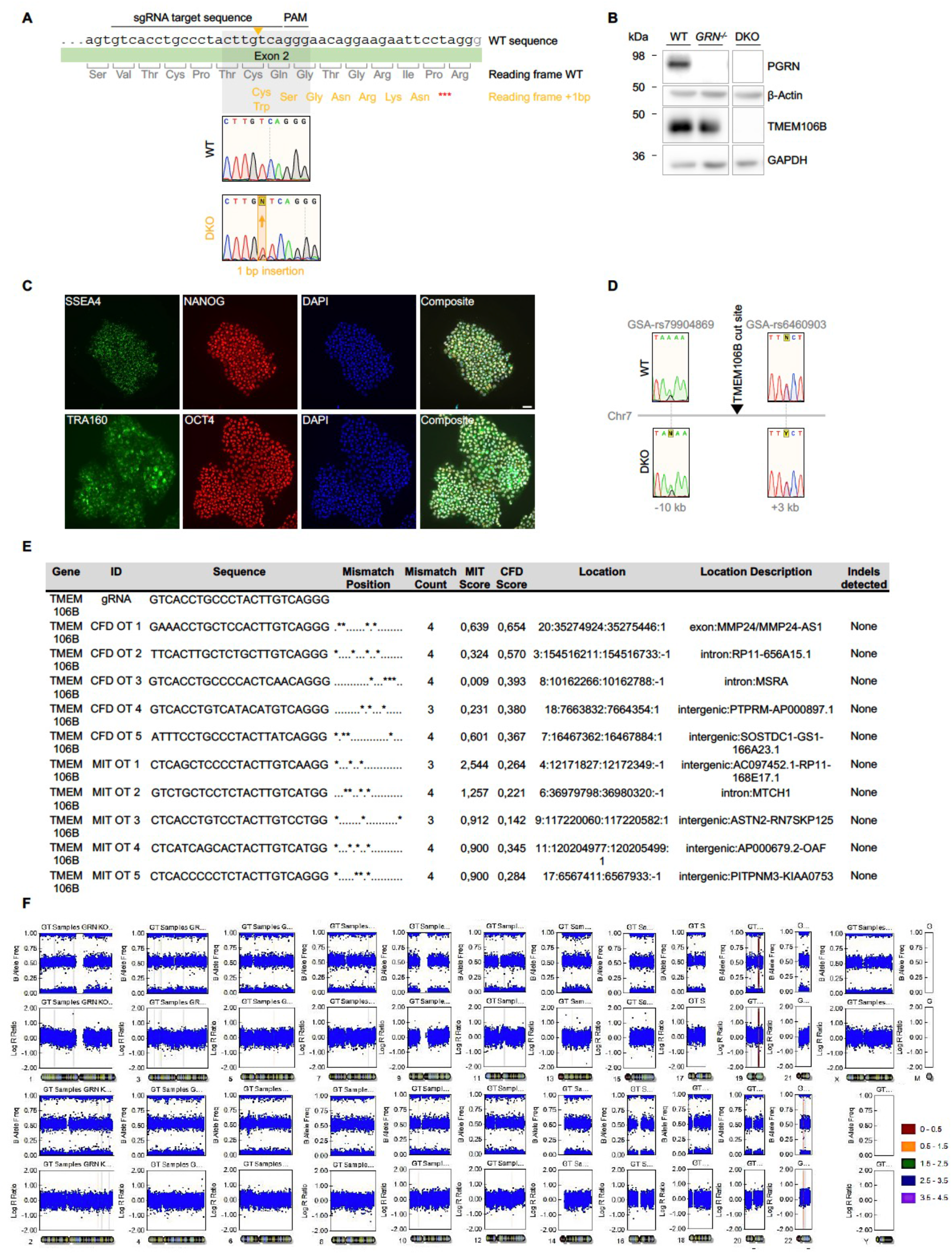
Generation of *TMEM106BxGRN* DKO hiPSC. (**A**) Schematic overview of the edited locus on exon 2 of the TMEM106B gene. Indicated are the sgRNA target sequence, the protospacer adjacent motif (PAM), reading frame with WT and DKO amino acid sequence and Sanger sequencing traces of WT and DKO lines. (**B**) Western blot confirming loss of TMEM106B and hPGRN in edited hiPSC in comparison to WT. (**C**) Immunofluorescence images showing expression of pluripotency markers SSEA4, NANOG, TRA160 and OCT4 (scalebar = 50 µm) in edited iPSCs. (**D**) Sanger sequencing of heterozygous SNPs on both sides of the edited locus in WT and DKO confirm that no loss of heterozygosity was induced by CRISPR editing. (**E**) Overview of the most likely off-targets as assessed using CRISPOR. Sanger sequencing of the PCR-amplified off-target sequences showed no deviation from the WT sequence. (**F**) Karyogram of DKO cell line indicating integrity of all chromosomes after editing, as assessed by molecular karyotyping.

**Table S1:**
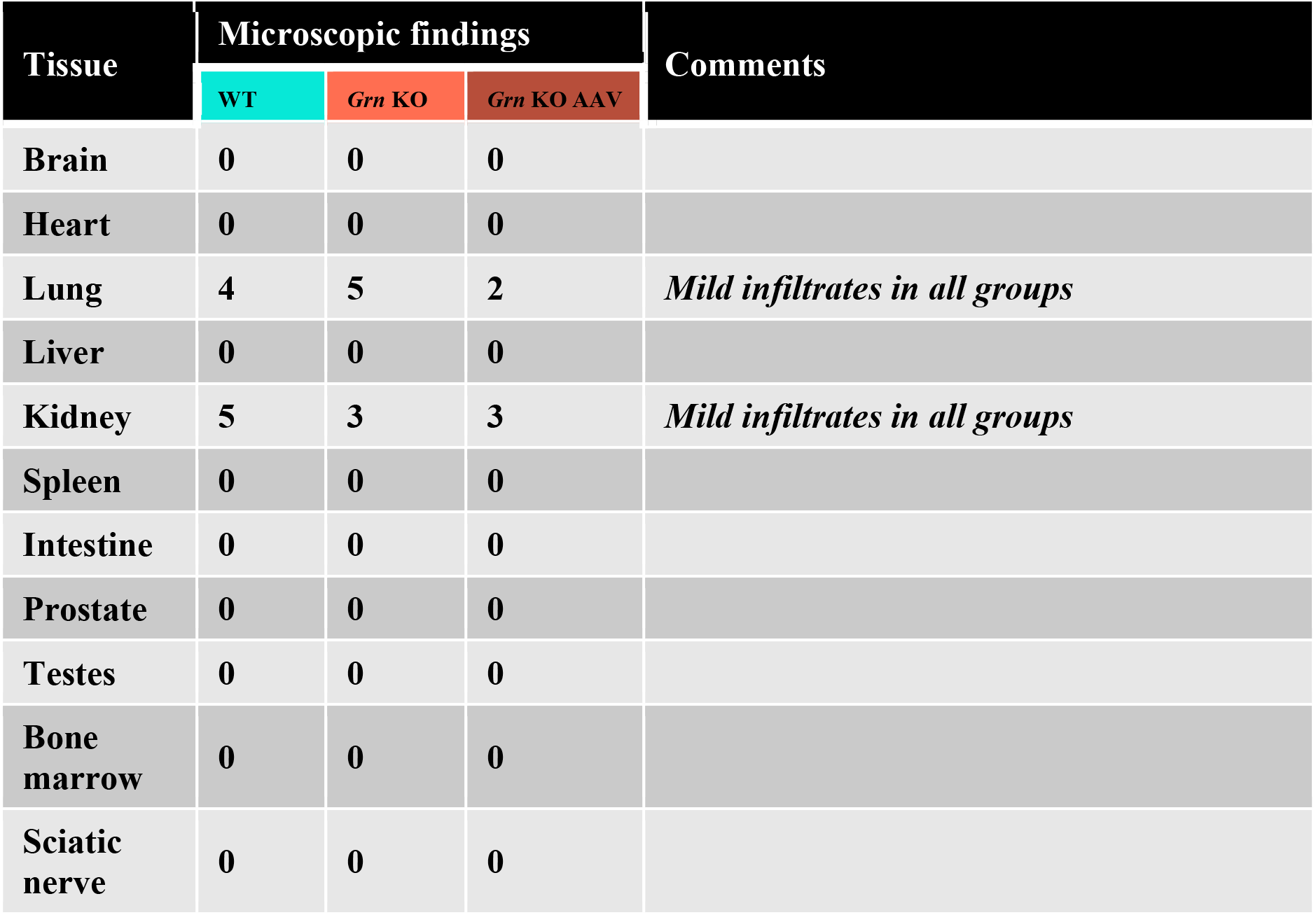
Summary of histopathological findings in Grn KO mice treated with AAV(L):bPGRN.

**Table S2:** Lipidomics in positive mode acquisition parameters.

| Name | Internal Standard | Q1 m/z | Q3 m/z | CE (V) |
| --- | --- | --- | --- | --- |
| 1-O-Palmitoyl-Cer(d18:1/18:0) | Cer(d18:1/16:0(d7)) | 786.8 | 502.5 | 35 |
| 24-Hydroxycholesterol | 24-Hydroxcholesterol-d7 | 385.3 | 367.3 | 30 |
| 24-Hydroxycholesterol(d7) | IS | 392.3 | 367.3 | 30 |
| 3-O-SulfoLacCer(d18:1/18:0) | LacCer(d18:1/17:0) | 970.8 | 548.5 | 61 |
| 4-beta-Hydroxycholesterol | Cholesterol(d7) | 420.3 | 385.3 | 15 |
| 7-keto-Cholesterol | Cholesterol(d7) | 401.3 | 383.3 | 15 |
| CE HETE | CE(18:1(d7)) | 706.6 | 369.2 | 25 |
| CE HODE | CE(18:1(d7)) | 682.6 | 369.2 | 25 |
| CE HpODE | CE(18:1(d7)) | 698.6 | 369.2 | 25 |
| CE oxoHETE | CE(18:1(d7)) | 704.6 | 369.2 | 25 |
| CE oxoODE | CE(18:1(d7)) | 680.6 | 369.2 | 25 |
| CE(16:1) | CE(18:1(d7)) | 640.6 | 369.3 | 26 |
| CE(18:1(d7)) | IS | 675.2 | 369.4 | 26 |
| CE(18:1) | CE(18:1(d7)) | 668.6 | 369.3 | 26 |
| CE(18:2) | CE(18:1(d7)) | 666.6 | 369.3 | 26 |
| CE(20:4) | CE(18:1(d7)) | 690.6 | 369.3 | 26 |
| CE(20:5) | CE(18:1(d7)) | 688.6 | 369.3 | 26 |
| CE(22:6) | CE(18:1(d7)) | 714.6 | 369.3 | 26 |
| Cer(d18:0/16:0) | Cer(d18:1/16:0(d7)) | 540.6 | 284.3 | 40 |
| Cer(d18:0/18:0) | Cer(d18:1/16:0(d7)) | 568.7 | 284.3 | 40 |
| Cer(d18:0/24:0) | Cer(d18:1/16:0(d7)) | 652.9 | 284.3 | 40 |
| Cer(d18:0/24:1) | Cer(d18:1/16:0(d7)) | 650.9 | 284.4 | 40 |
| Cer(d18:1/16:0(d7)) | IS | 545.5 | 271.4 | 40 |
| Cer(d18:1/16:0) | Cer(d18:1/16:0(d7)) | 538.5 | 264.3 | 40 |
| Cer(d18:1/18:0) | Cer(d18:1/16:0(d7)) | 566.6 | 264.3 | 40 |
| Cer(d18:1/24:0) | Cer(d18:1/16:0(d7)) | 650.6 | 264.3 | 40 |
| Cer(d18:1/24:1) | Cer(d18:1/16:0(d7)) | 648.6 | 264.3 | 40 |
| Cholesterol | Cholesterol(d7) | 369.3 | 369.3 | 10 |
| Cholesterol(d7) | IS | 376.2 | 376.2 | 10 |
| Cholesteryl hexoside | CE(18:1(d7)) | 566.6 | 369.3 | 17 |
| Coenzyme Q10 | TG(15:0/18:1(d7)/15:0) | 863.3 | 197.2 | 35 |
| DG(15:0/18:1(d7)) | IS | 605.6 | 346.5 | 30 |
| DG(16:0_18:1) | DG(15:0/18:1(d7)) | 612.4 | 313.3 | 30 |
| DG(16:0_20:4) | DG(15:0/18:1(d7)) | 634.5 | 313.3 | 30 |
| DG(18:0_18:1) | DG(15:0/18:1(d7)) | 640.4 | 341.3 | 30 |
| DG(18:0_20:4) | DG(15:0/18:1(d7)) | 662.5 | 341.3 | 30 |
| DG(18:0_22:6) | DG(15:0/18:1(d7)) | 686.6 | 341.3 | 30 |
| DG(18:1_20:4) | DG(15:0/18:1(d7)) | 660.5 | 339.3 | 30 |
| DG(18:1/18:1) | DG(15:0/18:1(d7)) | 638.4 | 339.3 | 30 |
| GB3(d18:1/16:0) | GB3(d18:1/18:0(d3)) | 1025 | 520.5 | 40 |
| GB3(d18:1/18:0(d3)) | IS | 1056 | 551.6 | 40 |
| GB3(d18:1/18:0) | GB3(d18:1/18:0(d3)) | 1053 | 548.6 | 40 |
| GB3(d18:1/24:0) | GB3(d18:1/18:0(d3)) | 1137 | 632.6 | 40 |
| GB3(d18:1/24:1) | GB3(d18:1/18:0(d3)) | 1135 | 630.6 | 40 |
| GlcCer(d18:1(d5)/18:0) | IS | 733.6 | 269.3 | 45 |
| GlcCer(d18:1/16:0(d3)) | IS | 703.7 | 264.3 | 51 |
| Glucosylsphingosine(d5) | IS | 467.2 | 269.3 | 16 |
| HexCer(d18:1/12:0) | GlcCer(d18:1(d5)/18:0) | 644.5 | 264.3 | 40 |
| HexCer(d18:1/16:0) | GlcCer(d18:1(d5)/18:0) | 700.6 | 264.3 | 40 |
| HexCer(d18:1/18:0) | GlcCer(d18:1(d5)/18:0) | 728.6 | 264.3 | 40 |
| HexCer(d18:1/22:0) | GlcCer(d18:1(d5)/18:0) | 784.7 | 264.4 | 40 |
| HexCer(d18:1/24:0) | GlcCer(d18:1(d5)/18:0) | 812.7 | 264.3 | 40 |
| HexCer(d18:1/24:1) | GlcCer(d18:1(d5)/18:0) | 810.7 | 264.3 | 40 |
| Hexosylsphingosine | Glucosylsphingosine(d5) | 462.3 | 264.2 | 16 |
| LacCer(d18:1/16:0) | LacCer(d18:1/17:0) | 862.6 | 264.3 | 40 |
| LacCer(d18:1/17:0) | IS | 876.6 | 264.3 | 40 |
| LacCer(d18:1/18:0) | LacCer(d18:1/17:0) | 890.7 | 264.3 | 40 |
| LacCer(d18:1/24:0) | LacCer(d18:1/17:0) | 974.8 | 264.3 | 40 |
| LacCer(d18:1/24:1) | LacCer(d18:1/17:0) | 972.7 | 264.3 | 40 |
| Lactosylsphingosine | Glucosylsphingosine(d5) | 624.4 | 264.3 | 16 |
| LPC(16:0) | LPC(18:1(d7)) | 496.3 | 184.1 | 40 |
| LPC(16:1) | LPC(18:1(d7)) | 494.5 | 184.1 | 40 |
| LPC(18:0) | LPC(18:1(d7)) | 524.3 | 184.1 | 40 |
| LPC(18:1(d7)) | IS | 529.3 | 184.1 | 40 |
| LPC(18:1) | LPC(18:1(d7)) | 522.3 | 184.1 | 40 |
| LPC(20:4) | LPC(18:1(d7)) | 544.3 | 184.1 | 40 |
| LPC(22:6) | LPC(18:1(d7)) | 568.3 | 184.1 | 40 |
| LPC(24:0) | LPC(18:1(d7)) | 608.5 | 184.1 | 40 |
| LPC(24:1) | LPC(18:1(d7)) | 606.5 | 184.1 | 40 |
| LPC(26:0) | LPC(18:1(d7)) | 636.5 | 104.1 | 40 |
| LPC(26:1) | LPC(18:1(d7)) | 634.5 | 104.1 | 40 |
| lyso-GB3 | lyso-GB3-d7 | 786.6 | 264.3 | 46 |
| lyso-GB3-d7 | IS | 793.5 | 271.3 | 46 |
| lyso-GB4 | GB3(d18:1/18:0(d3)) | 990.6 | 264.3 | 52 |
| MG(16:0) | MG(18:1(d7)) | 348.3 | 239.3 | 22 |
| MG(16:1) | MG(18:1(d7)) | 346.3 | 237.3 | 22 |
| MG(18:0) | MG(18:1(d7)) | 376.3 | 267.3 | 22 |
| MG(18:1(d7)) | IS | 381.3 | 272.5 | 22 |
| MG(18:1) | MG(18:1(d7)) | 374.3 | 265.3 | 22 |
| MG(20:4) | MG(18:1(d7)) | 396.3 | 287.3 | 22 |
| N-Oleoylethanolamine | LPC(18:1(d7)) | 326.3 | 62.1 | 23 |
| N-Palmitoyl-O-phosphocholineserine | LPC(18:1(d7)) | 509.5 | 184.1 | 40 |
| Palmitoylethanolamine | LPC(18:1(d7)) | 300.3 | 62.1 | 23 |
| PC(15:0/18:1(d7)) | IS | 754.6 | 184.1 | 40 |
| PC(16:0/5:0(CHO)) | PC(15:0/18:1(d7)) | 594.5 | 184.1 | 40 |
| PC(16:0/9:0(CHO)) | PC(15:0/18:1(d7)) | 650.4 | 184.1 | 40 |
| PC(16:0/9:0(COOH)) | PC(15:0/18:1(d7)) | 666.4 | 184.1 | 40 |
| PC(18:0/20:4(OH[S])) | PC(15:0/18:1(d7)) | 826.6 | 184.1 | 40 |
| PC(18:0/20:4(OOH[S])) | PC(15:0/18:1(d7)) | 842.6 | 184.1 | 40 |
| PC(34:1) | PC(15:0/18:1(d7)) | 760.6 | 184.1 | 40 |
| PC(36:1) | PC(15:0/18:1(d7)) | 788.6 | 184.1 | 40 |
| PC(36:2) | PC(15:0/18:1(d7)) | 786.6 | 184.1 | 40 |
| PC(36:4) | PC(15:0/18:1(d7)) | 782.6 | 184.1 | 40 |
| PC(38:4) | PC(15:0/18:1(d7)) | 810.6 | 184.1 | 40 |
| PC(38:6) | PC(15:0/18:1(d7)) | 806.6 | 184.1 | 40 |
| PC(40:5) | PC(15:0/18:1(d7)) | 836.6 | 184.1 | 40 |
| PC(40:6) | PC(15:0/18:1(d7)) | 834.6 | 184.1 | 40 |
| PC(O-16:0/0:0) | LPC(18:1(d7)) | 482.3 | 184.1 | 40 |
| PC(O-16:0/2:0) | LPC(18:1(d7)) | 524.3 | 184.2 | 40 |
| PC(O-18:0/2:0) | LPC(18:1(d7)) | 552.5 | 184.1 | 40 |
| PE(15:0/18:1(d7)) | IS | 711.6 | 570.5 | 40 |
| PE(18:0/20:4(OH[S])) | PE(15:0/18:1(d7)) | 784.5 | 643.4 | 40 |
| PE(18:0/20:4(OOH[S])) | PE(15:0/18:1(d7)) | 800.5 | 659.4 | 40 |
| PE(34:1) | PE(15:0/18:1(d7)) | 718.6 | 577.6 | 40 |
| PE(36:1) | PE(15:0/18:1(d7)) | 746.6 | 605.5 | 40 |
| PE(36:2) | PE(15:0/18:1(d7)) | 744.6 | 603.5 | 40 |
| PE(36:4) | PE(15:0/18:1(d7)) | 740.6 | 599.5 | 40 |
| PE(38:4) | PE(15:0/18:1(d7)) | 768.6 | 627.5 | 40 |
| PE(38:5) | PE(15:0/18:1(d7)) | 766.6 | 625.5 | 40 |
| PE(38:6) | PE(15:0/18:1(d7)) | 764.6 | 623.5 | 40 |
| PE(40:4) | PE(15:0/18:1(d7)) | 796.6 | 655.5 | 40 |
| PE(40:5) | PE(15:0/18:1(d7)) | 794.6 | 635.5 | 40 |
| PE(40:6) | PE(15:0/18:1(d7)) | 792.6 | 651.5 | 40 |
| PE(40:7) | PE(15:0/18:1(d7)) | 790.6 | 649.5 | 40 |
| Sitosteryl hexoside | CE(18:1(d7)) | 594.6 | 397.4 | 17 |
| SM(d18:1(d9)/18:1) | IS | 738.7 | 184.1 | 40 |
| SM(d18:1/16:0) | SM(d18:1(d9)/18:1) | 703.6 | 184.1 | 40 |
| SM(d18:1/18:0) | SM(d18:1(d9)/18:1) | 731.6 | 184.1 | 40 |
| SM(d18:1/24:0) | SM(d18:1(d9)/18:1) | 815.7 | 184.1 | 40 |
| SM(d18:1/24:1) | SM(d18:1(d9)/18:1) | 813.7 | 184.1 | 40 |
| Sphinganine | Sphingosine(d17:1) | 302.2 | 284.3 | 20 |
| Sphinganine 1-phosphate | Sphingosine 1-phosphate-d7 | 382.3 | 284.3 | 18 |
| Sphingosine | Sphingosine(d17:1) | 300.2 | 264.3 | 20 |
| Sphingosine 1-phosphate | Sphingosine 1-phosphate-d7 | 380.3 | 264.3 | 25 |
| Sphingosine 1-phosphate-d7 | IS | 387.3 | 271.3 | 25 |
| Sphingosine 1-phosphocholine | LPC(18:1(d7)) | 465.5 | 184.1 | 40 |
| Sphingosine(d17:1) | IS | 286.2 | 250.3 | 20 |
| TG(15:0/18:1(d7)/15:0) | IS | 829.8 | 570.8 | 40 |
| TG(18:0_36:2) | TG(15:0/18:1(d7)/15:0) | 904.7 | 603.4 | 40 |
| TG(18:1_34:2) | TG(15:0/18:1(d7)/15:0) | 874.7 | 575.4 | 40 |
| TG(18:1_34:3) | TG(15:0/18:1(d7)/15:0) | 872.7 | 573.4 | 40 |
| TG(20:4_32:1) | TG(15:0/18:1(d7)/15:0) | 870.6 | 549.3 | 40 |
| TG(20:4_34:2) | TG(15:0/18:1(d7)/15:0) | 896.6 | 575.3 | 40 |
| TG(20:4_34:3) | TG(15:0/18:1(d7)/15:0) | 894.6 | 573.3 | 40 |
| TG(20:4_36:0) | TG(15:0/18:1(d7)/15:0) | 928.8 | 607.5 | 40 |
| TG(20:4_36:2) | TG(15:0/18:1(d7)/15:0) | 924.7 | 603.4 | 40 |
| TG(20:4_36:3) | TG(15:0/18:1(d7)/15:0) | 922.7 | 601.4 | 40 |
| TG(22:6_36:2) | TG(15:0/18:1(d7)/15:0) | 948.7 | 603.4 | 40 |
| TG(22:6_38:1) | TG(15:0/18:1(d7)/15:0) | 978.7 | 633.4 | 40 |
| TG(22:6_38:2) | TG(15:0/18:1(d7)/15:0) | 976.7 | 631.4 | 40 |

Note: IS - Internal standard

**Table S3:**
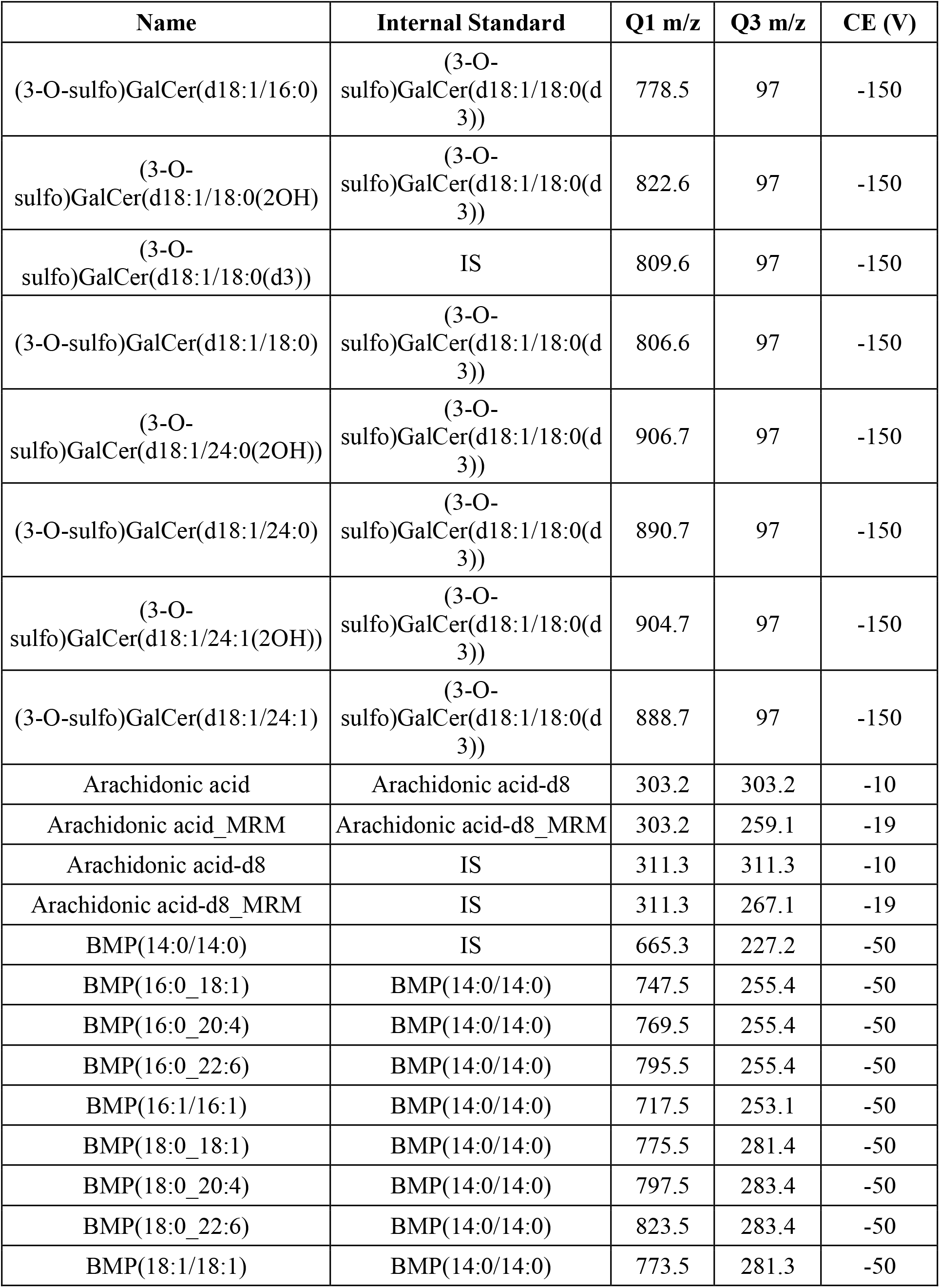
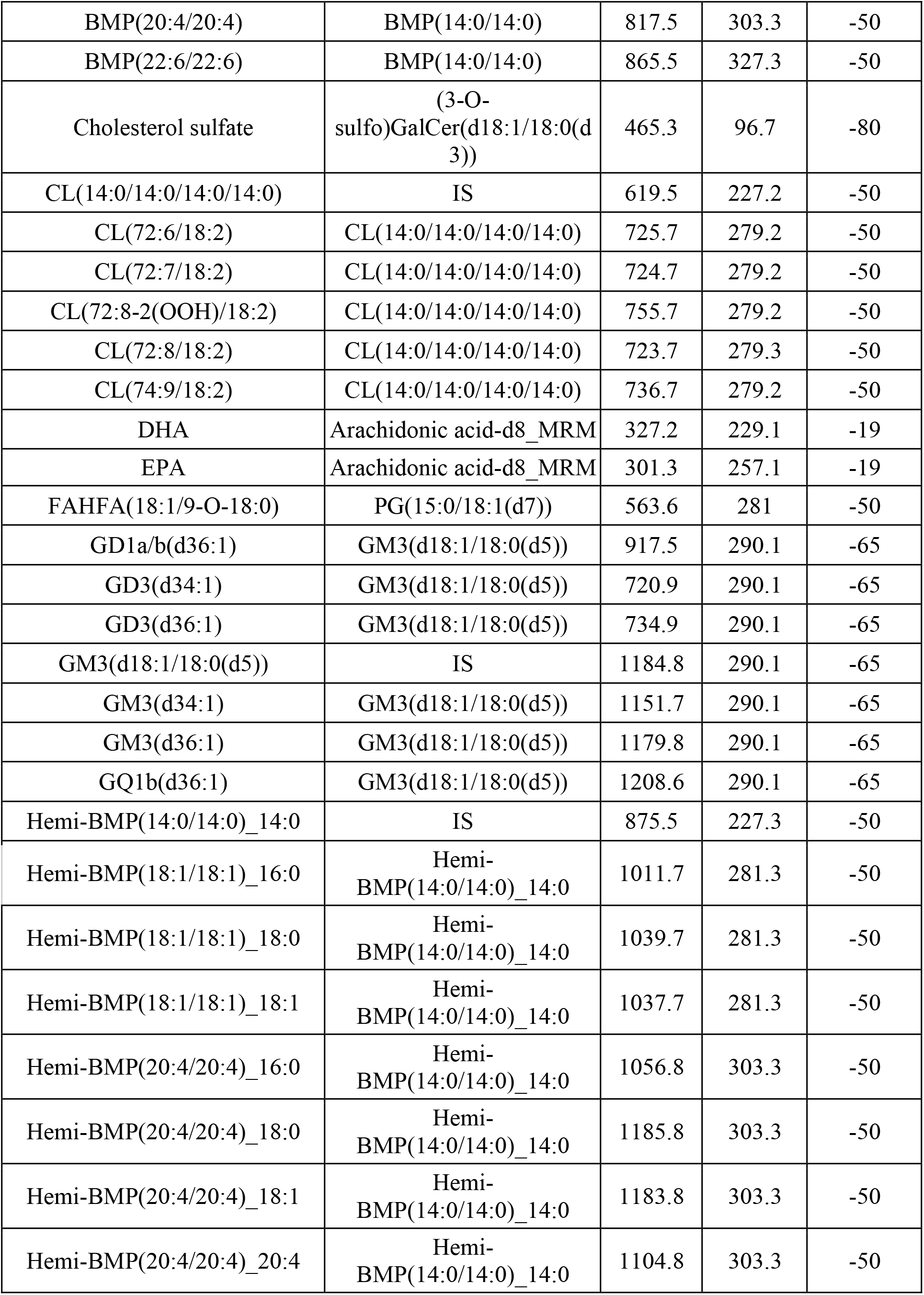
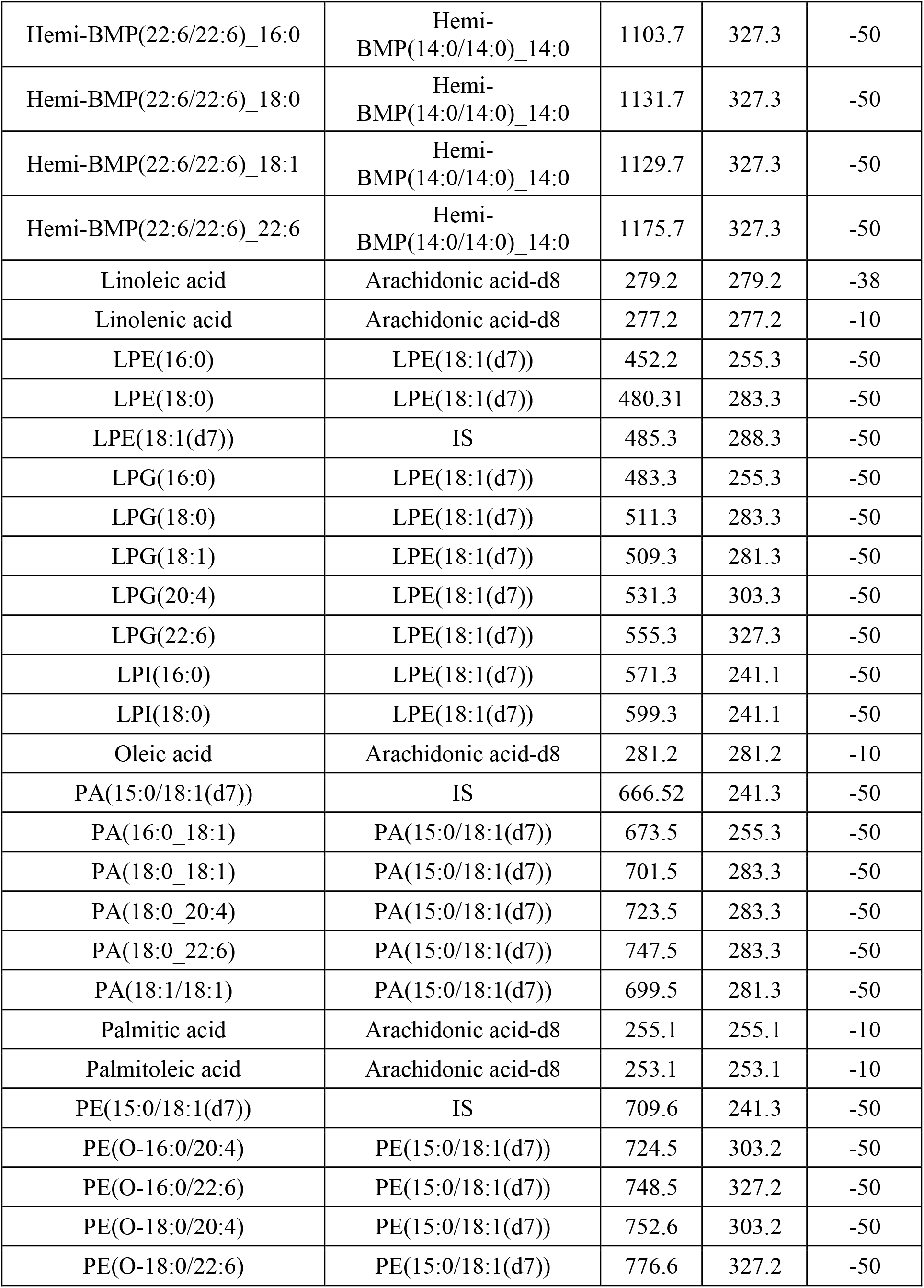
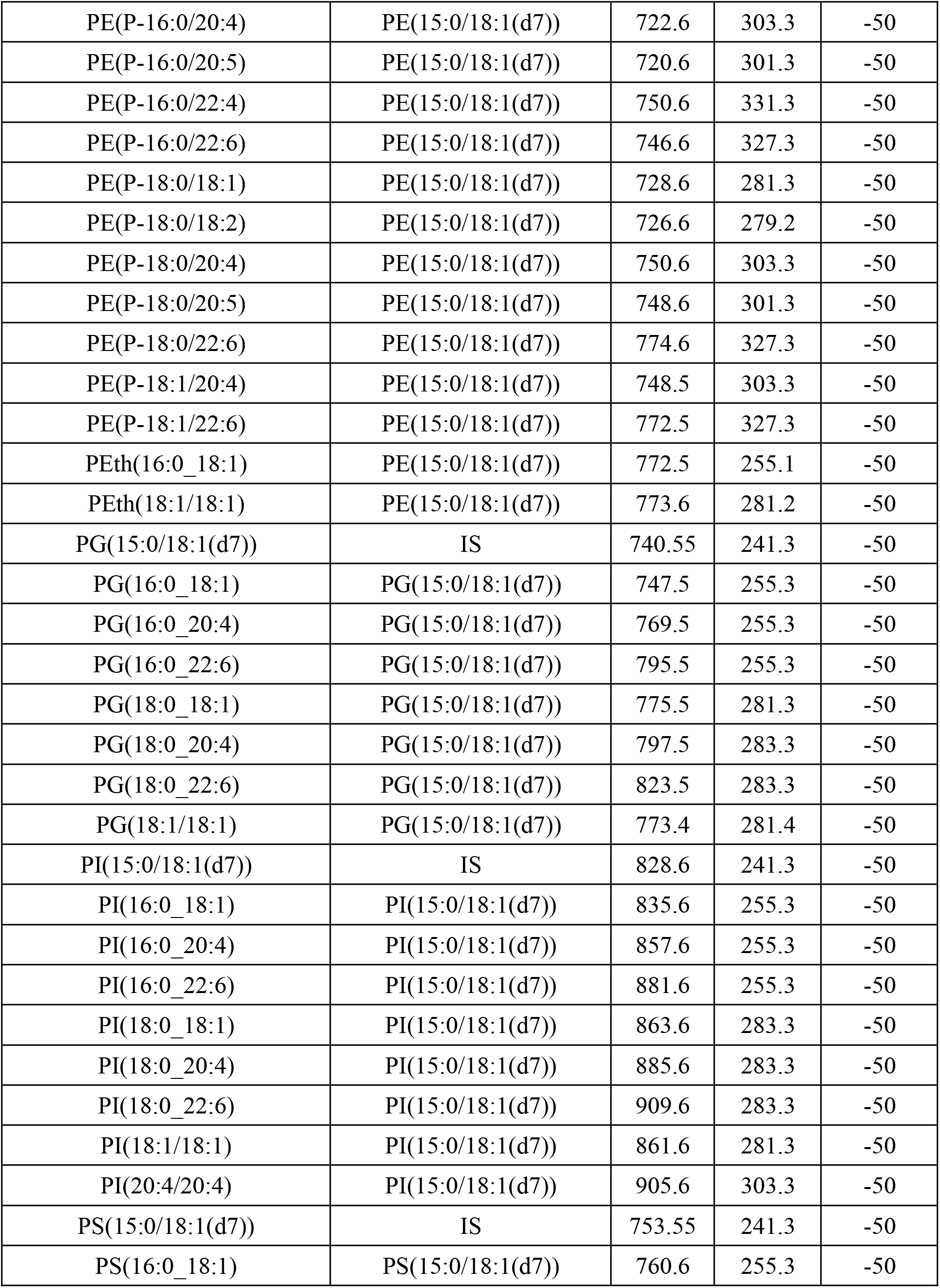
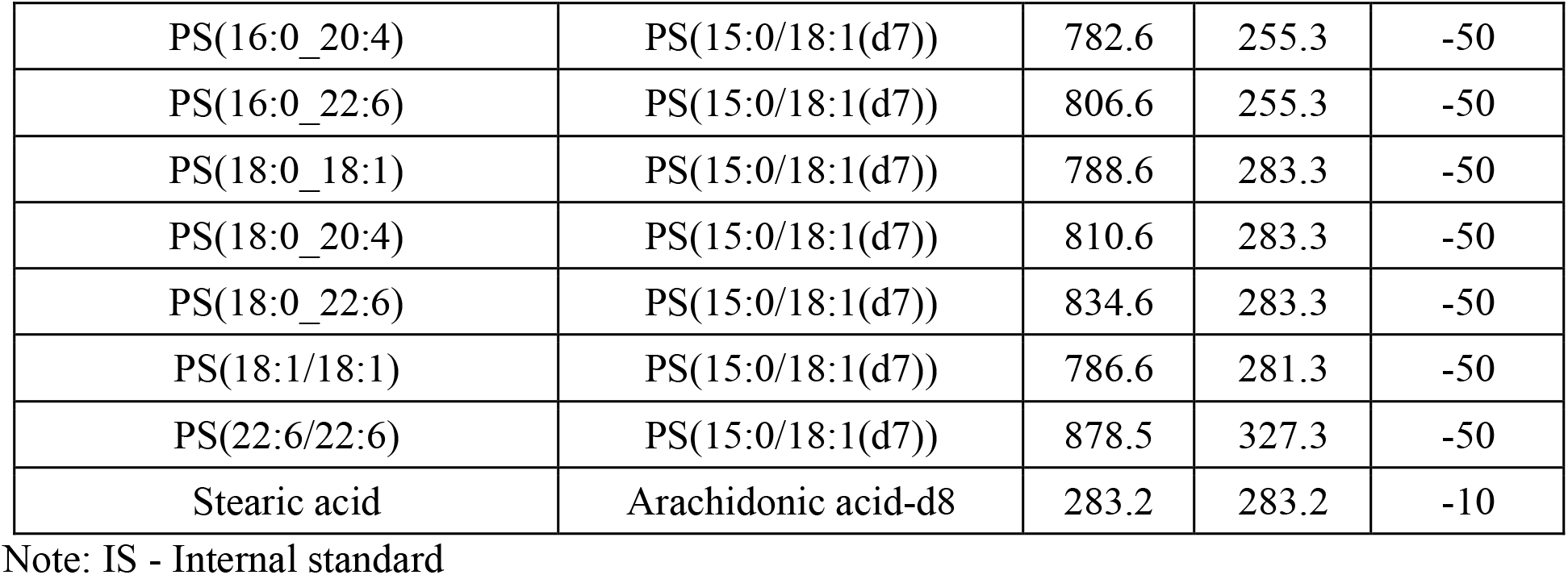
Lipidomics in negative mode acquisition parameters.

**Table S4:** Glucosylsphingolipid in positive mode acquisition parameters.

| Name | Internal Standard | Q1 m/z | Q3 m/z | CE (V) |
| --- | --- | --- | --- | --- |
| alpha-GalCer(d18:1/16:0) | GalCer(d18:1/15:0) | 700.6 | 264.3 | 32 |
| alpha-GalCer(d18:1/18:0) | GalCer(d18:1/15:0) | 728.6 | 264.3 | 32 |
| alpha-GalCer(d18:1/20:0) | GalCer(d18:1/15:0) | 756.6 | 264.3 | 32 |
| alpha-GalCer(d18:1/22:0) | GalCer(d18:1/15:0) | 784.7 | 264.3 | 32 |
| alpha-GalCer(d18:1/22:1) | GalCer(d18:1/15:0) | 782.7 | 264.3 | 32 |
| alpha-GalCer(d18:1/24:0) | GalCer(d18:1/15:0) | 812.7 | 264.3 | 32 |
| alpha-GalCer(d18:1/24:1) | GalCer(d18:1/15:0) | 810.7 | 264.3 | 32 |
| alpha-GalCer(d18:2/18:0) | GalCer(d18:1/15:0) | 726.6 | 262.3 | 32 |
| alpha-GalCer(d18:2/20:0) | GalCer(d18:1/15:0) | 754.6 | 262.3 | 32 |
| alpha-GalCer(d18:2/22:0) | GalCer(d18:1/15:0) | 782.7 | 262.3 | 32 |
| Cholesteryl galactoside | GlcCer(d18:1(d5)/18:0) | 566.6 | 369.3 | 13 |
| Cholesteryl glucoside | GlcCer(d18:1(d5)/18:0) | 566.6 | 369.3 | 13 |
| Galactosylsphingosine | Glucosylsphingosine(d5) | 462.2 | 282.3 | 18 |
| GalCer(d18:1/15:0) | IS | 686.3 | 264.3 | 32 |
| GalCer(d18:1/16:0) | GalCer(d18:1/15:0) | 700.6 | 264.3 | 32 |
| GalCer(d18:1/18:0) | GalCer(d18:1/15:0) | 728.6 | 264.3 | 32 |
| GalCer(d18:1/20:0) | GalCer(d18:1/15:0) | 756.6 | 264.3 | 32 |
| GalCer(d18:1/22:0) | GalCer(d18:1/15:0) | 784.7 | 264.3 | 32 |
| GalCer(d18:1/22:1) | GalCer(d18:1/15:0) | 782.7 | 264.3 | 32 |
| GalCer(d18:1/24:0) | GalCer(d18:1/15:0) | 812.7 | 264.3 | 32 |
| GalCer(d18:1/24:1) | GalCer(d18:1/15:0) | 810.7 | 264.3 | 32 |
| GalCer(d18:2/18:0) | GalCer(d18:1/15:0) | 726.6 | 262.3 | 32 |
| GalCer(d18:2/20:0) | GalCer(d18:1/15:0) | 754.6 | 262.3 | 32 |
| GalCer(d18:2/22:0) | GalCer(d18:1/15:0) | 782.7 | 262.3 | 32 |
| GlcCer(d18:1(d5)/18:0) | IS | 733.6 | 269.3 | 32 |
| GlcCer(d18:1/16:0(d3)) | IS | 703.6 | 264.3 | 32 |
| GlcCer(d18:1/16:0) | GlcCer(d18:1/16:0(d3)) | 700.6 | 264.3 | 32 |
| GlcCer(d18:1/18:0) | GlcCer(d18:1(d5)/18:0) | 728.6 | 264.3 | 32 |
| GlcCer(d18:1/20:0) | GlcCer(d18:1(d5)/18:0) | 756.6 | 264.3 | 32 |
| GlcCer(d18:1/22:0) | GlcCer(d18:1(d5)/18:0) | 784.7 | 264.3 | 32 |
| GlcCer(d18:1/22:1) | GlcCer(d18:1(d5)/18:0) | 782.7 | 264.3 | 32 |
| GlcCer(d18:1/24:0) | GlcCer(d18:1(d5)/18:0) | 812.7 | 264.3 | 32 |
| GlcCer(d18:1/24:1) | GlcCer(d18:1(d5)/18:0) | 810.7 | 264.3 | 32 |
| GlcCer(d18:2/18:0) | GlcCer(d18:1(d5)/18:0) | 726.6 | 262.3 | 32 |
| GlcCer(d18:2/20:0) | GlcCer(d18:1(d5)/18:0) | 754.6 | 262.3 | 32 |
| GlcCer(d18:2/22:0) | GlcCer(d18:1(d5)/18:0) | 782.7 | 262.3 | 32 |
| Glucosylsphingosine | Glucosylsphingosine(d5) | 462.2 | 282.3 | 18 |
| Glucosylsphingosine(d5) | IS | 467.2 | 287.3 | 18 |
| Sitosteryl galactoside | GlcCer(d18:1(d5)/18:0) | 594.6 | 397.4 | 13 |
| Sitosteryl glucoside | GlcCer(d18:1(d5)/18:0) | 594.6 | 397.4 | 13 |

Note: IS - Internal standard

Extra files:

MovieS1_flipping_test_WT.mov

MovieS2_flipping_test_WT_AAV.mov

MovieS3_ flipping_test_DKO.mov

MovieS4_flipping_test_DKO_AAV.mov

Fig4_RNAseq_data.xlsx

Fig5_LCMS_data.xlsx

FigS5_RNAseq_data.xlsx

FigS7_LCMS_data.xslx

